# Exportin-1 functions as an adaptor for transcription factor-mediated docking of chromatin at the nuclear pore complex

**DOI:** 10.1101/2024.05.09.593355

**Authors:** Tiffany Ge, Donna Garvey Brickner, Kara Zehr, D. Jake VanBelzen, Wenzhu Zhang, Christopher Caffalette, Sara Ungerleider, Nikita Marcou, Brian Chait, Michael P. Rout, Jason H. Brickner

## Abstract

Nuclear pore proteins (Nups) in yeast, flies and mammals physically interact with hundreds or thousands of chromosomal sites, which impacts transcriptional regulation. In budding yeast, transcription factors mediate interaction of Nups with enhancers of highly active genes. To define the molecular basis of this mechanism, we exploited a separation-of-function mutation in the Gcn4 transcription factor that blocks its interaction with the nuclear pore complex (NPC) without altering its DNA binding or activation domains. SILAC mass spectrometry revealed that this mutation reduces the interaction of Gcn4 with the highly conserved nuclear export factor Crm1/Xpo1. Crm1 both interacts with the same sites as Nups genome-wide and is required for Nup2 to interact with the yeast genome. *In vivo*, Crm1 undergoes extensive and stable interactions with the NPC. *In vitro*, Crm1 binds to Gcn4 and these proteins form a complex with the nuclear pore protein Nup2. Importantly, the interaction between Crm1 and Gcn4 does not require Ran-GTP, suggesting that it is not through the nuclear export sequence binding site. Finally, Crm1 stimulates DNA binding by Gcn4, supporting a model in which allosteric coupling between Crm1 binding and DNA binding permits docking of transcription factor-bound enhancers at the NPC.

## Introduction

Nuclear pore complexes (NPCs) are large proteinaceous channels spanning the nuclear envelope that mediate trafficking of macromolecules between the cytoplasm and the nucleus. These eightfold symmetrical ring structures are made up of 16-48 copies of ∼30 different pore proteins called nucleoporins (Nups) ^1–3^. In addition to this transport function, NPCs also physically associate with and regulate the genome. Specifically, Nups have been shown to interact with hundreds to thousands of sites in the genome in budding yeast, flies and mammals ^4–9^ . Whereas in yeast, this interaction with Nups correlates with localization of chromatin at the nuclear periphery, whereas in flies and mammals, Nups interact with chromatin both at the periphery and in the nucleoplasm ^8,10^. Nups promote diverse functions: stronger transcription, DNA repair, chromosome folding, gene silencing, and epigenetic transcriptional poising ^4,11–19^. However, the molecular mechanism by which particular Nups are recruited to specific sites in the genome is not well understood.

An excellent model for Nup-genome interactions is the recruitment of inducible genes to the NPC upon activation in budding yeast ^11,20^. Many inducible genes reposition to the nuclear periphery and physically associate with Nups upon activation. Previous work found that localization to the nuclear periphery requires several Nups, including the nucleoplasmic basket proteins Nup1, Nup2 and Nup60 ^12,13,19,21^. Furthermore, although localization to the nuclear periphery does not require transcription itself ^12,22^, peripheral localization is controlled by transcription factor (TF) binding sites ^13,19,23,24^. These TF binding sites function as *DNA zip codes*; when inserted at an ectopic site in the genome, they are sufficient to induce peripheral localization and interaction with Nups ^13,19,23^. Loss of the TFs or the zip codes disrupts targeting to the nuclear periphery and results in a quantitative decrease in transcription ^13,25^. Nucleoporins also promote transcription in flies, plants, and mammals^9,10,26,27^.

A number of yeast TFs have been shown to mediate peripheral localization: Cbf1, Gcn4, Put3, Sfl1, and Ste12 ^23,24,28^. This is likely a common function of yeast TFs; a screen of yeast DNA binding proteins found that ∼65% of them, when tethered to an ectopic locus, were sufficient to promote Nup2-dependent repositioning to the nuclear periphery ^29^. In mammalian cells, Nups interact strongly with super-enhancers that are rich in TF binding sites ^30^. Thus, mediating interaction with the NPC may be a widespread (but not universal) function of TFs.

How do TFs mediate interaction with Nups? Structure-function analysis of Gcn4, a TF induced during amino acid starvation, identified a well-conserved 27-amino acid fragment (aa 205-231) that is sufficient to mediate peripheral localization ^29^. This Positioning Domain (PD_GCN4_) is within the central domain of Gcn4, which is separate from its activation and DNA binding domains ^31^. Point mutations in the PD_GCN4_ disrupted the ability of Gcn4 to position its target genes to the nuclear periphery and resulted in lower levels of transcription. Thus, the *gcn4-pd* mutation represents a separation-of-function allele that perturbs Gcn4-mediated peripheral localization without altering its other functions.

To better define the molecular mechanism of TF-mediated targeting to the NPC, we performed quantitative proteomics of proteins that co-precipitate with wild type and *pd* mutant Gcn4. This experiment identified the major nuclear export factor Crm1/Xpo1 as enriched in the wild type sample. Crm1 inhibition or depletion led to rapid loss of peripheral localization and Nup-association of both Gcn4 target genes and non-Gcn4 target genes. This suggests that Crm1 is generally required for NPC-chromatin association and functions upstream of Nups. Furthermore, consistent with a previous study ^7^, we find that Crm1 interacts upstream of hundreds of highly transcribed genes, strongly overlapping and correlating with sites that interact with Nup1, Nup2 and Nup60. Either inhibiting Crm1 or depleting Nup2 led to a strong, global decrease in nascent transcription. Furthermore, Nup2 showed genetic interactions with subunits of the Mediator head and middle modules. Crm1 purified from yeast cell extracts copurifies with the entire NPC and shows preferential interaction with Nup2. Recombinant Crm1 interacts with Gcn4 through the PD_GCN4_ and DNA binding domain and this complex binds to Nup2 *in vitro*. Finally, the DNA binding activity of Gcn4 is stimulated by Crm1, suggesting allosteric coupling between DNA binding and protein binding. We conclude that Crm1 is a critical adaptor for TF-mediated docking of DNA at the NPC in yeast and may have a similar function in other organisms.

## Results

### Crm1/Xpo1 is required for localization of genes at the nuclear periphery

Mutations in the Gcn4 positioning domain disrupt Gcn4-mediated peripheral localization of *HIS4*. To confirm that this is true for other Gcn4 targets, an array of 128 lac operator binding sites (LacO array) was introduced downstream of four Gcn4 target genes, *HIS1*, *HIS2*, *HIS3*, *HIS5* and at the *URA3* locus bearing a single copy of the Gcn4 binding site (*URA3:Gcn4BS*). GFP-tagged Lac repressor and a nuclear envelope membrane protein were expressed in these cells and the colocalization of each locus with the nuclear envelope was measured using confocal microscopy ^11,32–34^. All of these loci repositioned to the nuclear periphery upon histidine starvation in the wild type strain but not in *gcn4-pd* strains (Figure 1A). Thus, mutations in the PD_GCN4_ disrupt peripheral localization of Gcn4 targets generally.

**Figure 1.**
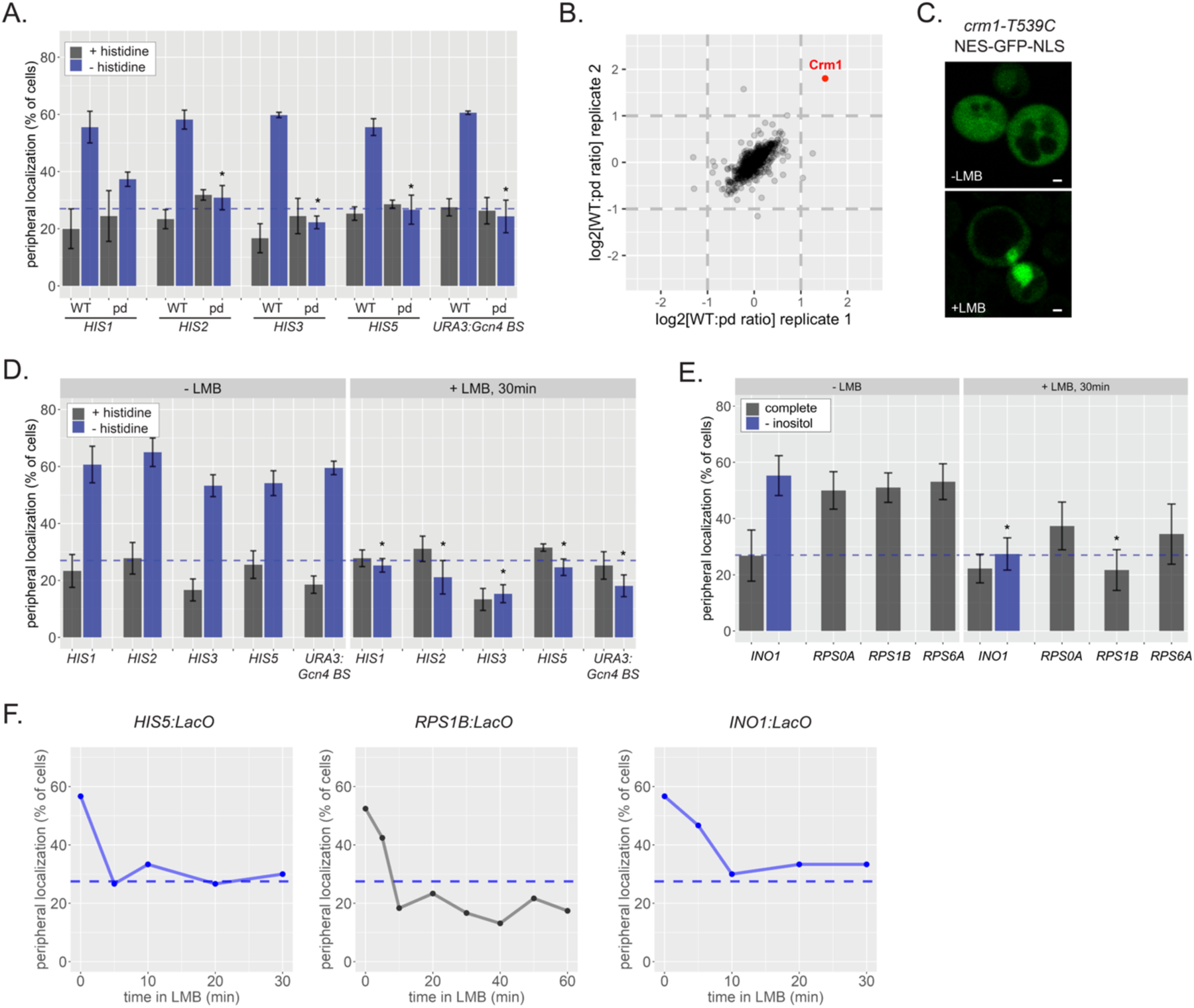
Crm1/Xpo1 is essential for peripheral targeting of genes that are both Gcn4-dependent and Gcn4-independent. **A.** Peripheral localization of four Gcn4 target genes the Gcn4 binding site inserted at an ectopic locus (*URA3:Gcn4BS*) in wild type or *gcn4-pd* mutant strains grown in SDC ± histidine. Mean percentage of the population ± standard error of the mean from ≥ three biological replicates of ≥ 30 cells each. Dashed line: localization expected for a randomly positioned gene. **B.** Scatter plot of the ratio of normalized abundance of 759 proteins identified by SILAC MS comparing the recovery from wild type (JBY558; light) and mutant (JBY557; heavy) cultures starved for histidine for 1h, with Crm1 highlighted. **C.** A mutant strain with a leptomycin B (LMB) sensitive allele of Crm1 (*crm1-T539C*) and expressing GFP bearing both a nuclear localization signal and a nuclear export signal (NES-GFP-NLS; untreated top panel) was treated with 100ng/ml LMB for 30 minutes (bottom). **D & E** Peripheral localization of either Gcn4 target genes (**D**) or non-Gcn4 target genes (**E**) in LMB-sensitive strains grown in in the indicated media treated ±100 ng/ml LMB for 30 minutes at 30°C. For panel E, only the *INO1:LacO* was grown in +inositol and -inositol medium. The *RPS* genes were localized in complete medium. **F.** Time courses of *HIS5*, *RPS1B* and *INO1* peripheral localization following addition of LMB.

To identify proteins that interact with Gcn4 in a PD_GCN4_-dependent manner, we immunopurified Gcn4-GFP from both wild type and *gcn4-pd* mutant strains grown in light and heavy lysine medium, respectively^35^. Gcn4-GFP was recovered from each lysate using anti-GFP nanobody-coupled magnetic beads (LaG16 ^36^). Associated proteins were pooled and proteins were identified by tandem mass spectrometry. Of 759 proteins identified (Supplementary Table 1), only Crm1/Xpo1 was significantly enriched (2-3-fold) in both technical replicates (Figure 1B). Thus, Crm1 shows PD_GCN4_-sensitive interactions with Gcn4.

Crm1 is a member of the karyopherin family of nucleocytoplasmic transport factors that interact with phe-gly (FG) repeat containing Nups in the NPC to mediate trafficking of cargoes ^37,38^; it is the major nuclear export factor (exportin) for proteins in yeast, raising the possibility that the positioning domain could function as a nuclear export sequence. However, we observed no difference in the nuclear-to-cytoplasmic ratio of wild type Gcn4-GFP compared with gcn4-pd-GFP (Figure S1). To test if Crm1 is important for Gcn4-mediated peripheral targeting, we utilized a yeast strain having the T539C mutation in Crm1, rendering it sensitive to the inhibitor leptomycin B (LMB ^39^), which binds covalently to the nuclear export sequence binding pocket of Crm1. The addition of LMB inhibited Crm1, leading to concentration of an NLS-GFP-NES reporter in the nucleus (Figure 1C). Into a *crm1-T539C* strain expressing GFP-LacI and an ER membrane marker, we introduced the LacO array at *HIS1*, *HIS2*, *HIS3, HIS5* and *URA3:Gcn4-BS* and expressed GFP-LacI and a nuclear envelope membrane marker ^34^. In the absence of LMB, these loci were targeted to the nuclear periphery normally upon histidine starvation (Figure 1D). However, in cells treated with LMB (100ng/ml, 30 minutes) these loci localized in the nucleoplasm instead. LMB had no effect on either the growth or the localization of these genes in yeast strains without the *crm1-T539C* mutation (not shown). Thus, inhibiting Crm1 disrupted peripheral localization of Gcn4 target genes.

To determine if Crm1 affects localization of non-Gcn4 targets, we tagged four other genes: the inducible gene *INO1*, which repositions to the nuclear periphery when cells are starved for inositol, or the constitutively expressed and peripheral genes *RPS0A*, *RPS1B* and *RPS6A* ^11,29^. Similar to the Gcn4 target genes, LMB treatment disrupted peripheral localization for all four of these non-Gcn4 targets (Figure 1E). Therefore, Crm1 is required for peripheral localization of genes that are both Gcn4-dependent and Gcn4-independent.

To assess how quickly LMB affects peripheral localization, we also examined the effect of LMB on the localization of *HIS5*, *RPS1B,* and *INO1* over time. In all three cases, the effect of LMB was very rapid; peripheral localization was disrupted within 5-10 minutes of LMB addition (Figure 1F). This is consistent with a direct role for Crm1 in mediating localization to the nuclear periphery.

### Crm1 and Nups interact upstream of hundreds of genes

Crm1 and Nups have been previously shown to interact with many sites in the yeast genome ^7^. However, because this study used microarrays for gene coding sequences that omitted intergenic regions, the precise location of these interactions was unclear. To define how Crm1 interacts with the yeast genome and to understand how its binding is related to the interaction of Nups, we employed chromatin endogenous cleavage sequencing (ChEC-seq2). Crm1, the nucleoplasmic basket proteins Nup1, Nup2 and Nup60 and the inner ring Nup157 were tagged with micrococcal nuclease, an extracellular endonuclease from *Staphylococcus aureus* ^40^. This non-specific endonuclease has an optimal calcium concentration of 10mM ^41^ and is inactive *in vivo*. MNase digestion can be stimulated by permeabilizing cells with detergent and adding 2mM Ca^+2^ and the cleavage sites can be identified by next generation sequencing ^42,43^. The cleavage pattern is then compared with cleavage by nuclear soluble MNase (sMNase) to control for non-specific cleavage of unprotected DNA.

The Crm1, Nup1, Nup2 and Nup60 MNase constructs cleaved the yeast genome significantly more than sMNase and had similar binding patterns genome-wide. At strongly transcribed genes such as *TDH3*, MNase-tagged Crm1, Nup1, Nup2 and Nup60 cleaved upstream of the promoter and within the upstream activating sequence region (i.e., the enhancers; Figure 2A). Cleavage by Nup157-MNase was quite weak and comparable to sMNase. Metagene plots generated by averaging the cleavage over all genes for each of these proteins highlighted that Crm1, Nup1, Nup2 and to some extent Nup60 cleave upstream of the promoter (Figure S2A). To inspect the types of genes that interact with Nups and Crm1 and the binding patterns of Crm1 and Nups at different gene subsets, we utilized several broad categories of genes defined by their association with transcription factors and co-regulators ^44^. These categories are: ribosomal protein genes (137 RPGs, very strongly expressed and co-regulated), genes that interact with coregulators such as the SAGA histone acetyltransferase, the Tup1 repressor or Mediator (1023 STM genes; typically strongly expressed, inducible genes), genes that interact with the pioneer factors Abf1 and Reb1 but do not recruit the STM coregulators (1549 TFO genes) and finally, genes that interact with RNA polymerase II but do not interact with known TFs or coregulators (2712 UNB genes; expressed at a lower level). The relative transcription level for each of these gene categories is RPGs > STMs > TFO > UNB (Figure S2B). Crm1, Nup1, Nup2 and Nup60 cleavage reflects transcription factor and coregulator binding (Figure 2B). Cleavage by these proteins was apparent upstream of the RPGs, the STM genes and the TFO genes, but not upstream of the UNB genes (Figure 2B). Nup157 cleavage was too low to detect, likely because it is farther from the genome.

**Figure 2.**
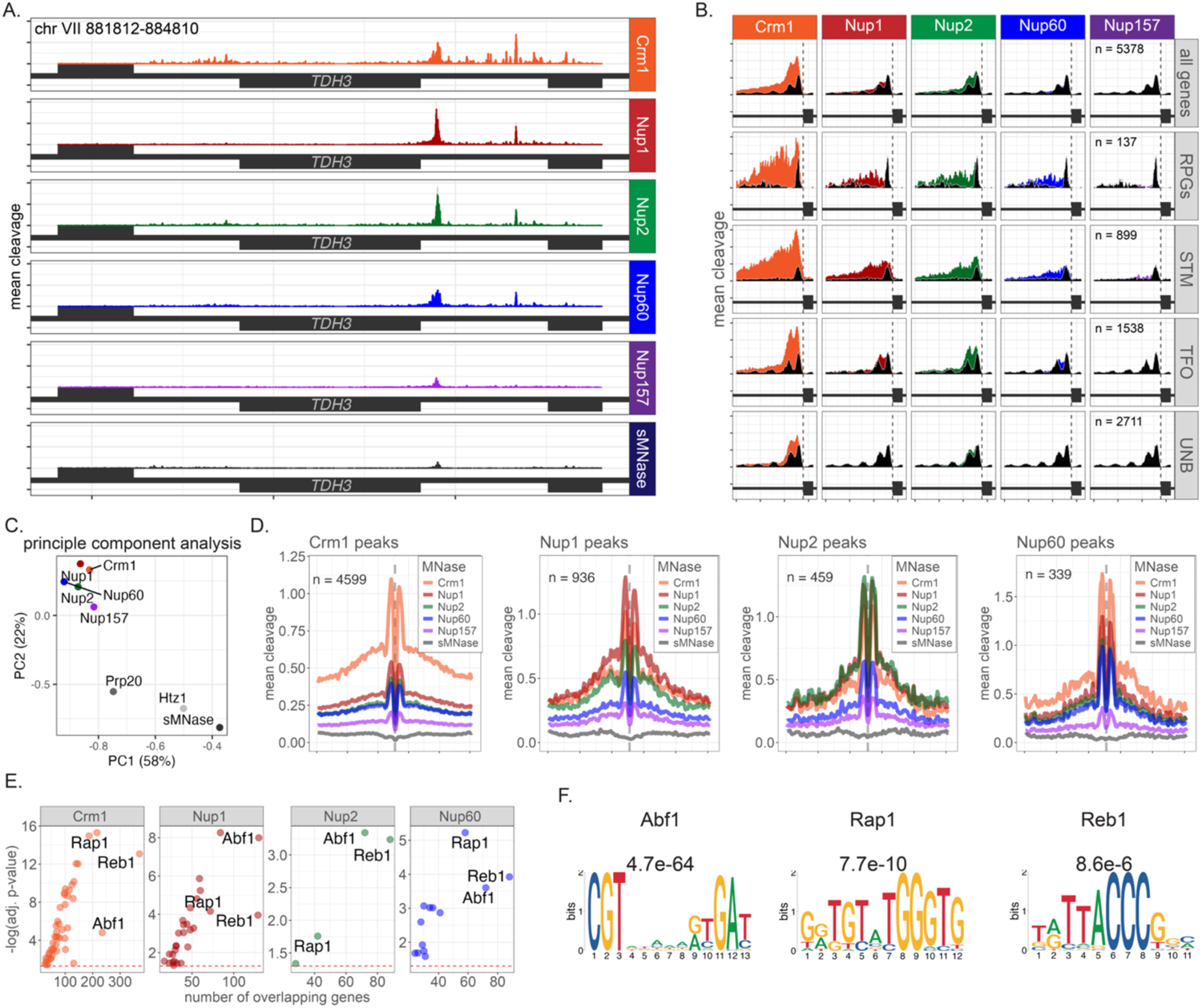
Crm1 and nuclear pore protein binding to the yeast genome. **A.** Mean CPM normalized ChEC cleavage frequency of Crm1-MNase, Nup1-MNase, Nup2-MNase, Nup60-MNase, Nup157-MNase, and soluble MNase near the *TDH3* gene. **B.** Metagene plots (normalized coding sequence length + 700 bp upstream and downstream) of the average CPM normalized ChEC cleavage over all RNAPII transcribed genes. Black plots = soluble MNase signal. **C.** Principal component analysis of cleavage frequency 700bp upstream of all genes by Crm1, Nup1, Nup60, Nup2, and Nup157 as well as controls (Prp20, sMNase and H2A.Z). **D.** Metagene plots for subsets of genes with distinct mechanisms of recruitment of RNAPII all genes: ribosomal protein genes (RPGs), inducible genes with TF-associated and SAGA-, TUP1-, and/or Mediator-associated promoters (STM), genes with TF-associated promoters lacking STM cofactors (TFO), and genes that recruit RNAPII without apparent TFs or cofactors (UNB). **E.** Average cleavage pattern around sites identified as high-confidence sites based on Crm1, Nup1, Nup2, or Nup60 ChEC-seq tracks ± 250bp flanking the site. **F.** Overlap between genes with cleavage peaks upstream for Nups and transcription factors based on ChIPexo. The number of genes that overlap between those identified by peaks of Crm1 and Nups with the indicated TFs was plotted against the Bonferroni-adjusted p-value (Fisher’s Exact test). **G.** Top three MEME results from Nup1 peaks and respective E-values.

When the cleavage frequency by these proteins over all upstream regions (-700 to -125 relative to start of coding sequence) was compared with each other and several negative controls (sMNase, MNase tagged nucleosome binding protein Prp20 and or histone H2A.Z ^45^) by principal component analysis, the Nups clustered with each other and Crm1, separate from the control proteins (Figure 2C). Despite the very weak cleavage from Nup157, it clustered with the other Nups and Crm1. Thus, these proteins interact with a common set of genes.

To assess if Crm1 and Nups are recruited to overlapping enhancers through interaction with TFs, we first identified high-confidence cleavage sites using DoubleChEC, a ChEC-seq analysis pipeline ^45^ that identifies high-confidence TF sites based on their relative enrichment over sMNase and their cleavage pattern (i.e., pairs of cleavage peaks flanking a protected region). DoubleChEC identified hundreds of high-confidence sites for each of the Nups and thousands for Crm1 (Figure 2D). Thus, Crm1 cleaves many more locations that the Nups. However, when the cleavage patterns of all of the proteins was plotted over high-confidence sites identified for each factor, we observed a high degree of cleavage of all of the sites by Crm1, Nup1, Nup2 and Nup60. Nup157 showed weak cleavage that had the same double peak pattern as the others and sMNase showed no cleavage (Figure 2D). Importantly, whereas all of the proteins cleaved with similar intensity over the peaks identified by Nup1, Nup2 and Nup60, the peaks identified by Crm1 showed much stronger cleavage by Crm1 than by any Nups. This suggests that Crm1 interacts not only at Nup binding sites but other sites in the genome as well.

The genes near the high-confidence sites identified by each of the Nups and Crm1 were also compared with genes bound to each of 78 different TFs from ChIP-exo experiments ^44^. Genes near high-confidence sites for each of the proteins showed strong overlap with the targets of a small set of TFs (Figure S2C). However, the targets of three TFs consistently showed strong overlap with all of genes near high-confidence Nup sites: Abf1, Reb1 and Rap1 (Figure 2E). Indeed, motif discovery of enriched DNA motifs from the high-confidence Nup1 sites identified the elements for each of these TFs (Figure 2F; consensus motifs for each in Figure S2D). These TFs bind many highly expressed genes, enriched for glycolysis and ribosomal protein genes (Figure S2E) consistent with the average cleavage for these Nups upstream of the RPG and STM genes.

### Crm1 functions upstream of Nup2 in gene recruitment to the nuclear periphery

Crm1 localizes both in the nucleoplasm and at the nuclear periphery, suggesting it could interact with chromatin in the nucleosplam. If Crm1 is an adaptor for TFs to interact with the nuclear pore complex, then it should function upstream of Nups. We tested if Crm1 is required for Nup2 association, by inhibiting or depleting Crm1 and measuring Nup2 binding to chromatin by ChEC. For comparison, we also tested if depletion of Nup2 affected Crm1 binding to chromatin.

We utilized three different conditional alleles: the LMB-sensitive allele of Crm1-T539C, and auxin-inducible degron (AID) alleles of both Nup2 and Crm1^46^. Nup2-AID was efficiently degraded upon addition of auxin (Supplementary Figure S3A), disrupting peripheral localization of *INO1*, *RPS0A*, *RPS1B* and *RPS6A*, as expected^29^ (Figure 3A). Attempts to construct a Crm1-AID strain were unsuccessful, perhaps because of inhibitory interactions between the AID tag and Crm1 (modeled in Figure S3B). Because Crm1-GFP strains are viable and GFP is not predicted to interact with Crm1 (Figure S3B), we developed an alternative strategy to deplete Crm1-GFP.Crm1-GFP was targeted for degradation by targeting the GFP for degradation using an inducible GFP-binding protein (GBP) fused to the miniAID tag ^47^ After inducing GBP-miniAID and adding auxin, we were able to deplete most of Crm1-GFP (Figure S3C) and strongly inhibit growth of the strains (Figure S3D; we refer to this system as grAID for **<u>gr</u>**een fluorescence protein-mediated **<u>a</u>**uxin-**i**nducible **<u>d</u>**egradation).

**Figure 3.**
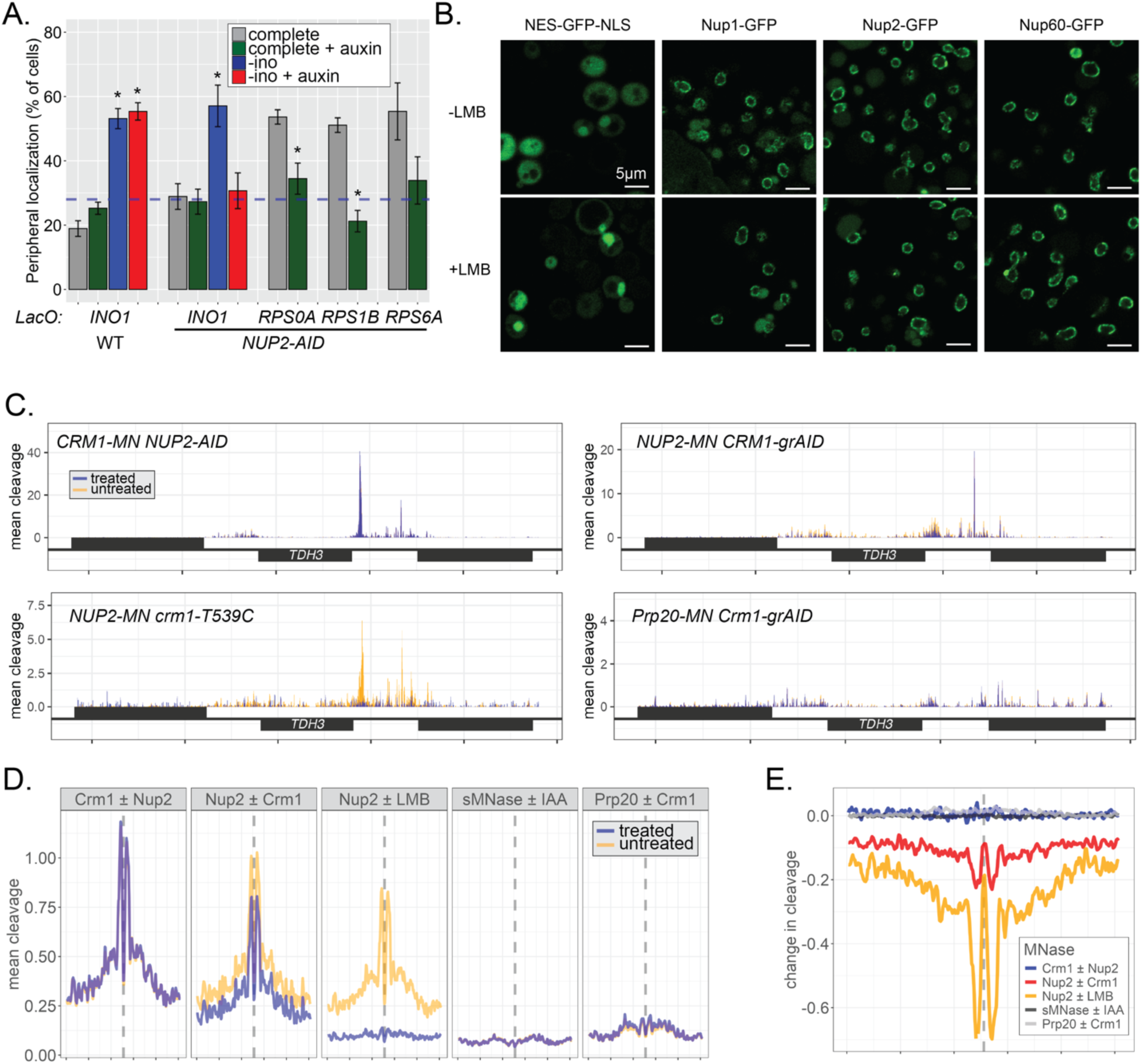
Crm1 functions upstream of Nup2. **A.** Localization of *INO1*, *RPS0A*, *RPS1B* and *RPS6A* in either a wild type strain (left) or a Nup2-AID strain ± auxin for 1h. Asterisks indicate p < 0.05, comparing to the untreated SDC control. **B.** Localization of GFP tagged proteins ± leptomycin B treatment. The NES-GFP-NLS reporter becomes concentrated in the nucleus, while Nup1, Nup2 and Nup60 are unaffected. **C - E.** Cleavage by Crm1, Nup2, Prp20 or sMNase in strains either depleted of Nup2 (by Nup2-AID), depleted of Crm1 (using grAID), or inhibited with leptomycin B. For this experiment, cleavage was carried out in SDC medium. Mean cleavage over the *TDH3* locus (**C**) or over Nup2 peaks (Figure 2E; panel **D**) from untreated or treated (i.e., +IAA (Nup2-AID), + 5-Ph-IAA and estradiol (Crm1-grAID) or +LMB (*crm1-T539C*). **E**. Difference in cleavage over Nup2 peaks between treated and untreated samples.

Crm1 ChEC cleavage over high-confidence Crm1 sites was unaffected by Nup2 depletion (Figure 3C & D). In contrast, either depleting Crm1 or inhibiting it with LMB led to decreased Nup2 cleavage upstream of highly expressed genes (Figure 3C) and over high-confidence Nup2 sites (Figure 3D). The control proteins sMNase and Prp20 were unaffected by either auxin treatment or Crm1 depletion (Figure 3C & D). LMB inhibition, while clearly blocking nuclear export, had no obvious effect on the localization of Nup1, Nup2 or Nup60 (Figure 3B). Therefore, Crm1 functions upstream of Nup2 and does not require Nup2 for its binding to chromatin. The effect of LMB was much stronger than the effect of Crm1 depletion, perhaps because the grAID system leads to incomplete depletion of Crm1-GFP (Figure S3C). Finally, since loss of Nup2 generally blocks targeting to the nuclear periphery, we surmise from the ChEC data that Crm1 still interacts with chromatin in the nucleoplasm.

### Crm1 and Nup2 promote transcription

Crm1 and Nups interact strongly with many highly expressed genes (Figure 2). In fact, ChEC with RNA polymerase II (Rpb1-MNase) revealed that the cleavage pattern of Rpb1-MNase upstream of all genes (700bp upstream to 25bp downstream of the transcription start site) was strongly correlated with that of Crm1 and Nups, with Spearman’s correlation values of 0.71-0.95 (Figure 4A). In contrast, the Rpb1-MNase cleavage pattern was poorly correlated with that of sMNase (correlation = 0.14).

**Figure 4.**
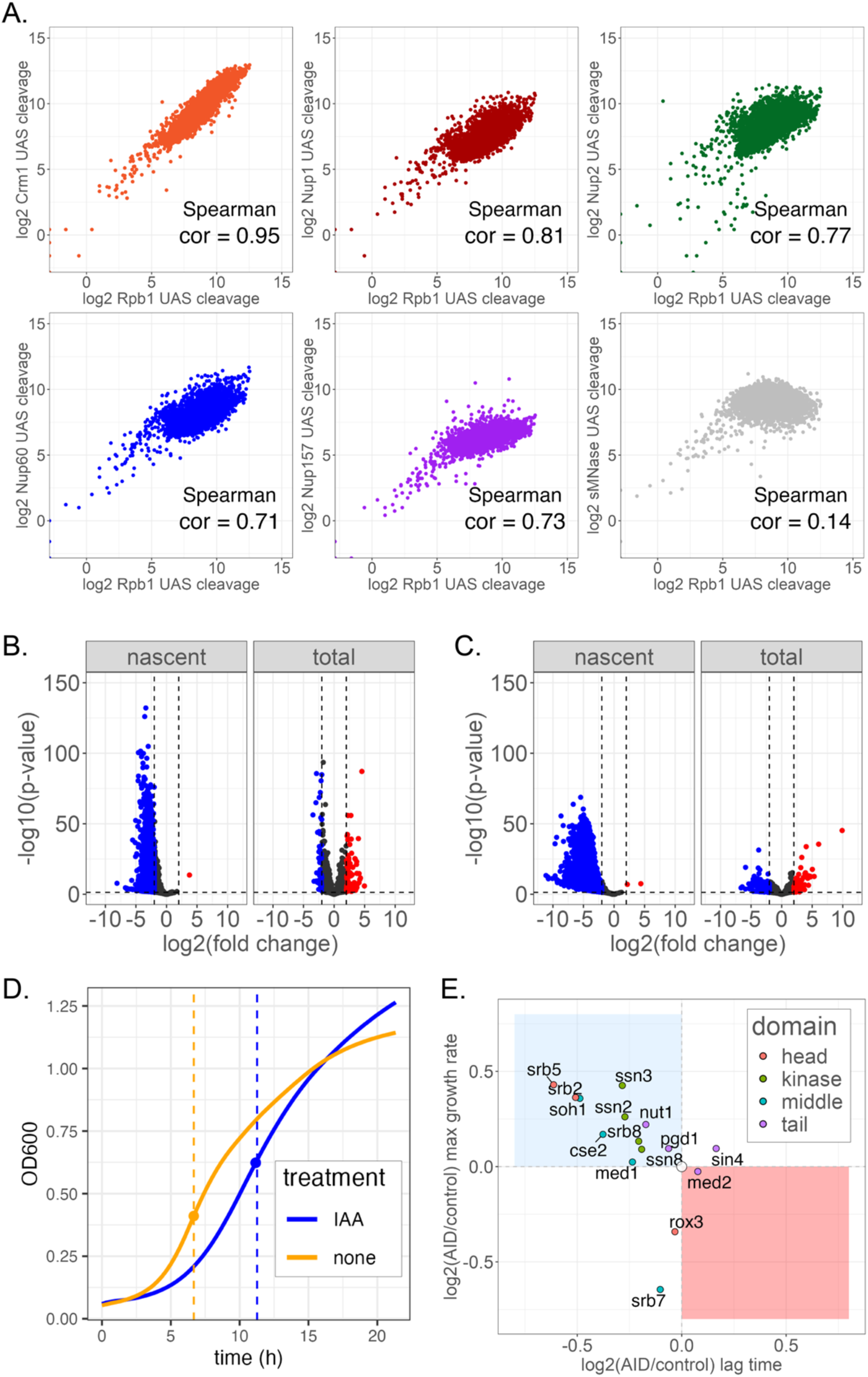
Crm1 and Nup2 promote stronger transcription. **A.** Scatter plots and Spearman’s correlation of Crm1 and Nup cleavage over upstream activating sequences (UASs; -700 to -125 from transcriptional start site) vs Rpb1-MNase cleavage for all yeast genes. **B & C**. SLAMseq analysis of nascent and total mRNA from either a *crm1-T539C* strain treated for 30 minutes with leptomycin B (**B**) or a Nup2-AID strain treated with auxin for 16h (**C**). **D**. Growth analysis of Nup2-AID strain ± auxin. The density of cultures of Nup2-AID strains was compared with Nup2-AID strains lacking non-essential Mediator subunits. The maximal growth rate (slope at the indicated timepoints) and the lag time (i.e., the time required to achieve the maximal growth rate) are highlighted. The effect of the loss of Nup2 was determined from the effects of adding auxin to the Nup2-AID strain. The fitness effect of loss of the Mediator mutations alone was determined from the ratio of these values to those of the Nup2-AID strain in the absence of auxin. The ratio of the maximal growth rate and lag time for each double mutant was compared with the expected value from multiplying the two growth defects (**E**). The blue square highlights buffering interactions and the red square highlights negative genetic interactions.

To determine if the interaction of Crm1 and nuclear pore proteins with enhancers affects transcription, we measured total and nascent RNA using thiol (SH)-linked alkylation for the metabolic sequencing of RNA (SLAM-seq^48^) following inhibition of Crm1 with LMB or depletion of Nup2 via the auxin-induced degradation (AID) system. Both inhibition of Crm1 (30 minutes; Figure 4B) and depletion of Nup2-AID (16h; Figure 4C) resulted in strong decreases in nascent mRNA levels. This supports a role for Crm1 and Nup2 in promoting stronger transcription.

While the inhibition of Crm1 and depletion of Nup2 resulted in strong downregulation in nascent transcription of hundreds of genes, this decrease was not reflected in the total mRNA levels (Figure 4B & C). This suggests that the defect associated with loss of Crm1 or Nups is strongly buffered through changes in mRNA half-life, a phenomenon that has been observed for numerous other perturbations that cause global defects in nascent transcription ^49–52^. Furthermore, the consequences of Nup2 depletion were not immediately apparent; after 1h of Nup2 depletion, we observed no significant effect on nascent transcript levels (data not shown). Thus, while the effect of Crm1 inhibition was immediate, the effects of Nup2 depletion were observed after several hours.

Previous work has suggested that disrupting the recruitment of genes to the nuclear periphery reduces the frequency of transcription ^25^, suggesting that loss of interaction with the NPC results in a defect in enhancer function ^53^. We explored the relationship between Crm1/Nups and Mediator function by quantitatively assessing genetic interactions between *NUP2* and each of the non-essential Mediator subunits. Null mutations for each of these subunits were introduced in Nup2-AID strains, all of which were viable (not shown). To see if Nup2 was genetically interacting with a particular subunit, we first measured the maximal growth rate (dashed line) ± auxin (Figure 4D, orange and blue dots, respectively) and the lag, or time required to reach that maximal growth rate (x intercept to dots in Figure 4D), for each strain. The fitness effect caused by loss of Nup2 is reflected in the increased lag and slower maximal growth rate in the presence of auxin (Figure 4D). The fitness effect of each of the Mediator mutants was measured in the *medΔ NUP2-AID* in the absence of auxin and fitness effect of loss of both Nup2 and a Mediator subunit was determined by measuring these parameters from the double mutant strains + auxin.

If the two mutations affect two unrelated biological functions, they should exhibit an additive fitness defect (white dot in Figure 4E), whereas, if the two mutations impact parallel, independent steps that contribute to the same process, then they should produce a larger-than-expected effect on fitness (i.e., negative genetic interaction; red quadrant in Figure 4E). Finally, if the two mutations impact the same step in a biological process, then loss of either will be equivalent to loss of both and they should show a less-than-additive “buffering” interaction (i.e., positive genetic interaction; blue quadrant in Figure 4E) ^54^. Most of the Mediator subunits exhibited buffering interactions with Nup2 (Figure 4E). Specifically, three of the four middle domain subunits (red dots), two of the three kinase domain subunits (green dots) and two of the three head subunits (teal dots) showed buffering interactions (highlighted in blue on the structure of Mediator, Figure S4). The tail subunits showed the weakest interactions and had either slightly positive or slightly negative interactions (purple dots in Figure 4E). Two genes, *ROX3* and *SRB7*, showed negative interaction on maximal growth rate, but simple additive effects on lag phase (Figure 4E; red in Figure S4). When mapped onto the structure of the Mediator complex, the buffering interactions and the negative interactions localize to distinct parts of Mediator (Figure S4). These data suggest that loss of Nup2 affects the same process as loss of the Mediator head, middle and kinase domains.

### Crm1 interacts stably with the NPC

To identify molecular interactions that may be relevant to targeting of genes to the nuclear periphery, we affinity-purified Crm1-GFP from yeast. Crm1 was tagged with GFP having a PreScission protease cleavage site upstream (Crm1-x-GFP) at the endogenous locus. Cells expressing this fusion protein were grown in rich medium, harvested, snap frozen and then lysed by cryomilling ^55^. Crm1-x-GFP was affinity purified using magnetic beads coupled to the LaG94-10 anti-GFP nanobody ^36^. A control lysate, lacking a GFP tagged protein, was subjected to the same purification. These affinity isolations resulted in recovery of Crm1-x-GFP and several strong co-purifying bands and very little background from the control lysate (Figure 5A, lanes 1 & 2). Furthermore, cleavage by PreScission protease resulted in elution of all of these co-purifying proteins (Figure 5A, lanes 3 & 4), suggesting that they are bound to the beads through Crm1. The eluted material was concentrated and subjected to trypsin digestion and tandem mass spectrometry, identifying 303 proteins (Supplementary Table S2). Label-free quantification (LFQ) of these proteins revealed all known subunits of the NPC, the paralog of Nup53, Asm4, and the transport factors Kap95, Srp1, Mex67, and Sac3 were among the top 42 hits (based on LFQ; Supplementary Table S2). Normalizing to the known stoichiometries of these proteins within the NPC ^56^ revealed a 10-fold range of intensities, due to Crm1 interacting with both the full isolated NPC but also with numerous different overlapping Nup subcomplexes ^57,58^ (Figure 5B). The strongest signals were from the RNA export platform (Nup159), the inner ring (Nsp1, Nup57 and Nup170, Nic96 and associated factors Nup116 and Gle2) and the nuclear basket FG repeat proteins Nup2 and Nup1. This suggest that Crm1 interacts stably with multiple parts of the NPC, including the nucleoplasmic face.

**Figure 5.**
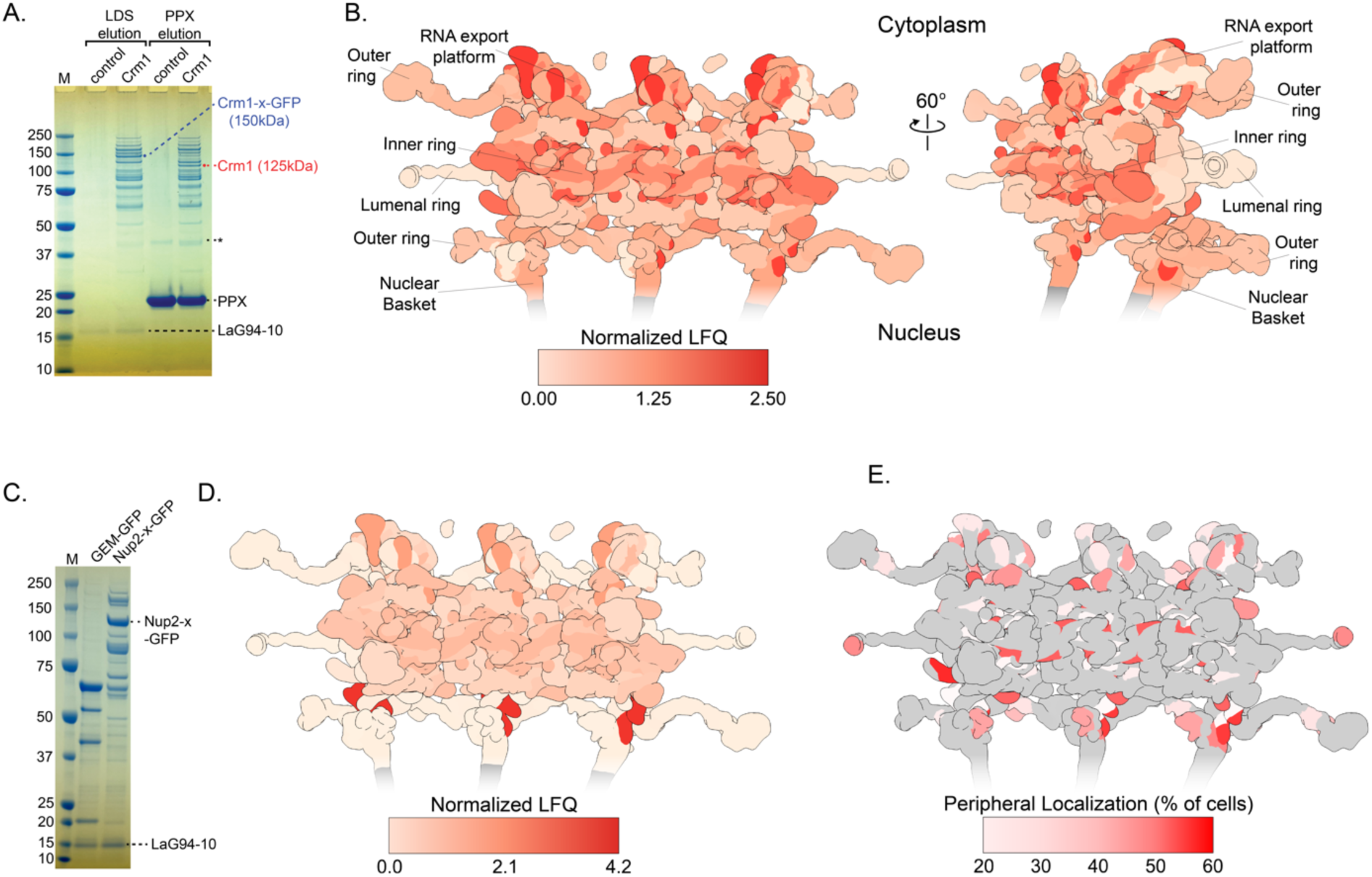
Crm1 and Nup2 interactions at the NPC overlap with the parts of the NPC that can contact chromatin. Crm1-x-GFP (panel **A**) or Nup2 (panel **C**) were purified from yeast using LaG94-10 anti-GFP nanobody magnetic beads. **A**. Lanes 2 and 4 were from the Crm1-x-GFP strain; lanes 1 and 3 are a mock purification from a lysate lacking a GFP tagged protein. For lanes 1 and 2, the proteins were eluted by heating in LDS sample buffer. For lanes 3 and 4, the proteins were eluted by PreScission protease (PPX) cleavage. Samples were subjected to mass spectrometry and label free quantification. **B & D**. The abundance of Nups from Crm1 (**B**) and Nup2 (**D**) was normalized to their stoichiometry within the pore to highlight over-represented hits. **C**. Lane 1 is a purification from a strain expressing Genetically Encoded Multimeric particles tagged with GFP ^85^; lane 2 is a purification from a strain expressing Nup2-x-GFP. Samples were eluted by heating in SDS and subjected to mass spectrometry and label free quantification. **D**. The 31 known nuclear pore proteins were identified in the top 224 hits. **E**. Heatmap of peripheral localization of *URA3* induced by fusing LexA to each of 27 Nups in a strain having a LacO array and LexA binding site at *URA3* (Figure S5).

For comparison, we also affinity purified Nup2-x-GFP (Figure 5C) and subjected the co-purifying proteins to tandem mass spectrometry. As with Crm1, we recovered all 32 known NPC proteins (Supplementary Table S2). However, normalizing to the stoichiometry of each protein within the pore revealed a 400-fold range of intensities, suggesting that Nup2 interacts directly with a much narrower range of Nups than Crm1. Consistent with previous studies ^59–63^, the most abundant protein from the Nup2 purification was Nup60 on the nucleoplasmic face of the NPC (Figure 5D). Proteins in the inner ring (Nsp1, Nup49, Nup53, Nup57, Nup170, Nup188, Nic96, Nup116 and Gle2) and the central channel GLFG protein Nup100 were ∼ 4-fold less than that of Nup60. Moreover, the outer ring, the cytoplasmic export complex, transmembrane proteins and the other nucleoplasmic proteins (Nup1, Mlp1 and Mlp2) were weakly recovered. Thus, Nup2 interacts most strongly with Nup60 in the nucleoplasmic face of the NPC and the inner ring, consistent with recent work ^59^.

We next asked which parts of the NPC can contact chromatin, to shed light on the connection between the chromatin binding of Nup2 and Crm1 and their association with the NPC. The LexA protein was fused to 27 nuclear pore proteins and these strains were crossed against a strain having a LacO array and the LexA binding site (LexA BS) at the *URA3* locus as well as the GFP-LacI and the ER/nuclear envelope membrane marker ^29^. *URA3* normally localizes in the nucleoplasm and we have used this strategy to test for the effect of tethering transcription factors on nuclear positioning ^23,29^. We reasoned that if LexA can come in contact with DNA, *URA3:LexABS* would localize at the nuclear periphery. Remarkably, over half of the Nups tested in this system were capable of repositioning *URA3:LexABS* to the nuclear periphery (Figure S5). Mapping these results onto the structure of the NPC, the Nups that were able to reposition chromatin to the nuclear periphery overlapped strongly with those that interact with Crm1 (Figure 5E). This suggests that the nucleoplasmic half of the NPC can contact chromatin, consistent with its open and flexible structure there.

### Biochemical reconstitution of Crm1 docking complex

To test the hypothesis that Crm1 serves as an adaptor for TF-NPC interactions, we asked if these proteins interact *in vitro*. First, to identify a minimal portion of Gcn4 that is sufficient to mediate peripheral localization of Gcn4 target genes, we expressed 100 amino acids of the carboxyl terminal portion of Gcn4 that includes both the PD and the DNA binding domain (PD-DBD). Expressing this fragment caused both *HIS4* and *HIS5* to reposition constitutively to the nuclear periphery (Figure 6A), suggesting that this fragment recapitulates Gcn4-mediated targeting *in vivo*.

**Figure 6.**
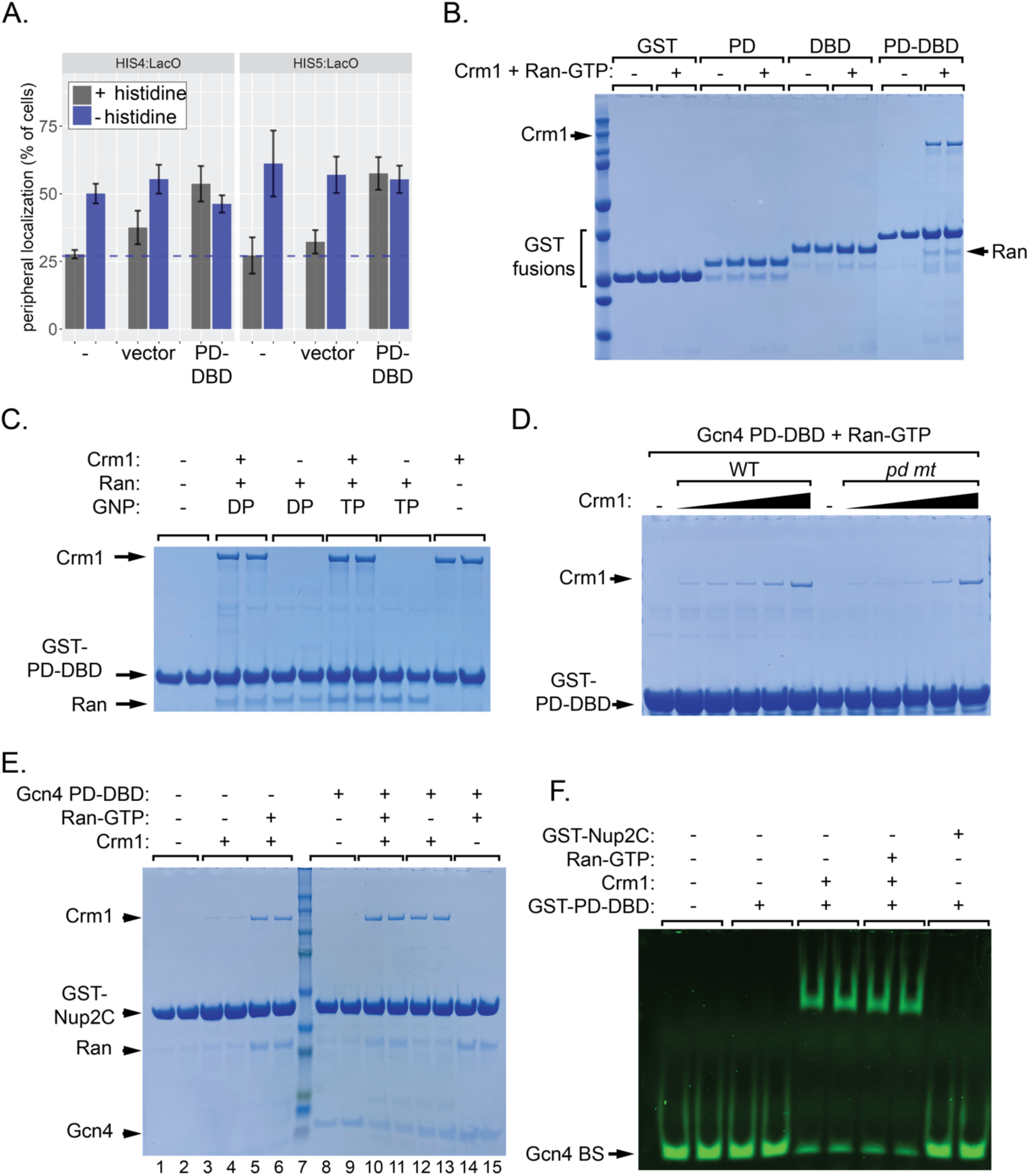
Biochemical reconstitution of TF-NPC bridging complex. **A**. *GCN4* strains transformed with the indicated constructs were grown in medium ± histidine and the localization of either *HIS4* or *HIS5* was determined. **B-E**. Recombinant His-tagged Crm1 (5µM) and His-tagged Gsp1/Ran (± GDP or GTP; 10µM) and untagged Gcn4 PD-DBD were incubated with the indicated GST fusion proteins on magnetic beads. Bead-bound proteins were washed twice, eluted with 40mM glutathione and separated by SDS-PAGE. The GST fusion proteins, Crm1 Gcn4 and Ran are indicated. **D**. Wild type or pd mutant GST-PD-DBD (amino acids 181-281) bound to magnetic beads were incubated with the following concentrations of recombinant Crm1: 338nM, 675nM, 1.35µM, 2.7µM and 6.75µM. **F**. 200nM Gcn4 DBD or PD-DBD were incubated with fluorescent Gcn4 binding site in the presence of the following concentrations of 1µM Crm1 ± 2.5µMRan-GTP.

Purified, recombinant GST fusion proteins bearing different fragments of Gcn4 on glutathione-conjugated magnetic beads were incubated with recombinant His_6_-tagged Crm1 and His_6_-tagged Ran (Gsp1) loaded with GTP. Crm1 and Ran bound to GST-PD-DBD and weakly to GST-DBD, but not to GST-PD or GST (Figure 6B). This suggests that the PD is not sufficient for interaction and that Crm1/Ran binds to the PD-DBD through both the PD and DBD domains. Indeed, Alphafold3 ^64^ structural predictions suggest that the PD folds back on the DNA binding domain of the opposite member of the dimer (Figure S6A).

Next, we tested if this interaction requires Ran-GTP, which is concentrated in the nucleus. Ran-GTP binding to Crm1 induces a conformational change that allows binding of nuclear export sequences (NESs) ^65^. Binding of Crm1 to GST-NES is strongly dependent on both Ran and GTP (Figure S6B). However, binding of Crm1 to Gcn4 PD-DBD is unaffected by either the nucleotide or Ran (Figure 6C). Fluorescence polarization shows that Crm1 alone binds fluorescent Gcn4 PD-DBD with a K_d_ ∼ 100nM (Figure S6C). Furthermore, the pd mutation leads to reduced binding of Crm1 at intermediate concentrations (Figure 6D), suggesting that this mutation reduces, rather than abolishing, binding. Therefore, Crm1 binding to Gcn4 PD-DBD is likely through a surface that is not affected by the conformational change induced by Ran-GTP.

Based on the mass spectrometry results and the requirement for Nup2 in peripheral localization, we hypothesized that Crm1-Gcn4 docks at the NPC through interaction with Nup2. Nup2 has a C-terminal Ran binding domain, which could interact with Crm1 through Ran-GTP, and twelve FxFG repeats that interact with Crm1 through its surface HEAT repeats ^66,67^. A similar complex has been reported between Crm1-Ran and Yrb2 ^67^. If so, then Ran should stimulate the interaction between Crm1 and Nup2. To test this prediction, we purified the C terminus of Nup2 (Nup2C; amino acids 496-721, including two FxFGs and the Ran binding domain) fused to GST. Both glutathione bead pulldowns (Figure 6E, lanes 3-6) and fluorescence polarization (Figure S6D) showed weak binding of Crm1 to GST-Nup2C that was strongly stimulated by Ran-GTP. Surprisingly, addition of Gcn4-PD-DBD also stimulated binding of Crm1 to Nup2C (Figure 6E, lanes 2 & 3 vs lanes 10 & 11; Figure S6D). Furthermore, we do not observe any apparent competition between Ran and Gcn4-PD-DBD, suggesting that all four proteins can bind simultaneously (Figure 6E).

Gcn4 is one of many TFs that mediate peripheral localization of genes. However, because these TFs do not show any obvious peripheral bias in their localization, it is unclear how they do this. We hypothesized that Crm1 and/or Nup2C interacts preferentially with DNA-bound TFs. If so, then allosteric coupling between Crm1 binding and DNA binding would lead the Crm1 interaction to stabilize the interaction between the TF and DNA. We tested this idea by adding Crm1 ± Ran-GTP or GST-Nup2C to Gcn4 in an electrophoretic mobility shift assay using fluorescently labeled Gcn4 binding site. For this experiment, we used a concentration of Gcn4 that results in shifting of a small fraction of the probe (Figure S6E). Upon addition of Crm1, with or without Ran-GTP, we observed a significant increase in the fraction of probe that was shifted (Figure 6F). This effect was not observed upon addition of Nup2C (Figure 6F). The mobility of the shifted species was the same as that observed in the absence of Crm1 (Figure S6E & data not shown), suggesting that this represents Gcn4-DNA complexes that do not contain Crm1. Nevertheless, this result suggests that Crm1 stimulates the DNA binding affinity of Gcn4.

## Discussion

The work presented here leads us to propose that Crm1/Xpo1 serves as an adaptor between TFs and nuclear pore proteins to both mediate chromatin-NPC interactions and promote transcription (Figure 7). This function is distinct from Crm1’s role as a nuclear export factor, for the interaction with Gcn4 does not require Ran-GTP. Thus, we anticipate that Gcn4 and likely other TFs bind to a surface on Crm1, likely one of its 21 HEAT repeats, that is not affected by the conformational change induced by Ran-GTP. Crm1 in the nucleoplasm interacts with DNA-bound TFs and this Crm1-TF-DNA complexes undergo random sub-diffusion until they encounter the NPC. At the NPC, Crm1 forms a complex with Nup2 (and potentially other Nups). Although the Crm1-Gcn4 interaction does not require Ran-GTP and the interaction of Crm1-Gcn4 with Nup2 are Ran-independent, it is possible that Ran-GTP can dissociate from the docking complex. However, the interaction of Nup2 with Nup60 on the NPC is enhanced by Ran-GTP ^61^, raising the possibility that Ran-GTP bound to Crm1 stabilizes the docking of Nup2-Crm1-Gcn4 on Nup60. Finally, the affinity of these interactions (∼100-200nM) is consistent with the transient, yet continuous targeting to the nuclear periphery that is observed in live cell tracking experiments ^68^.

**Figure 7.**
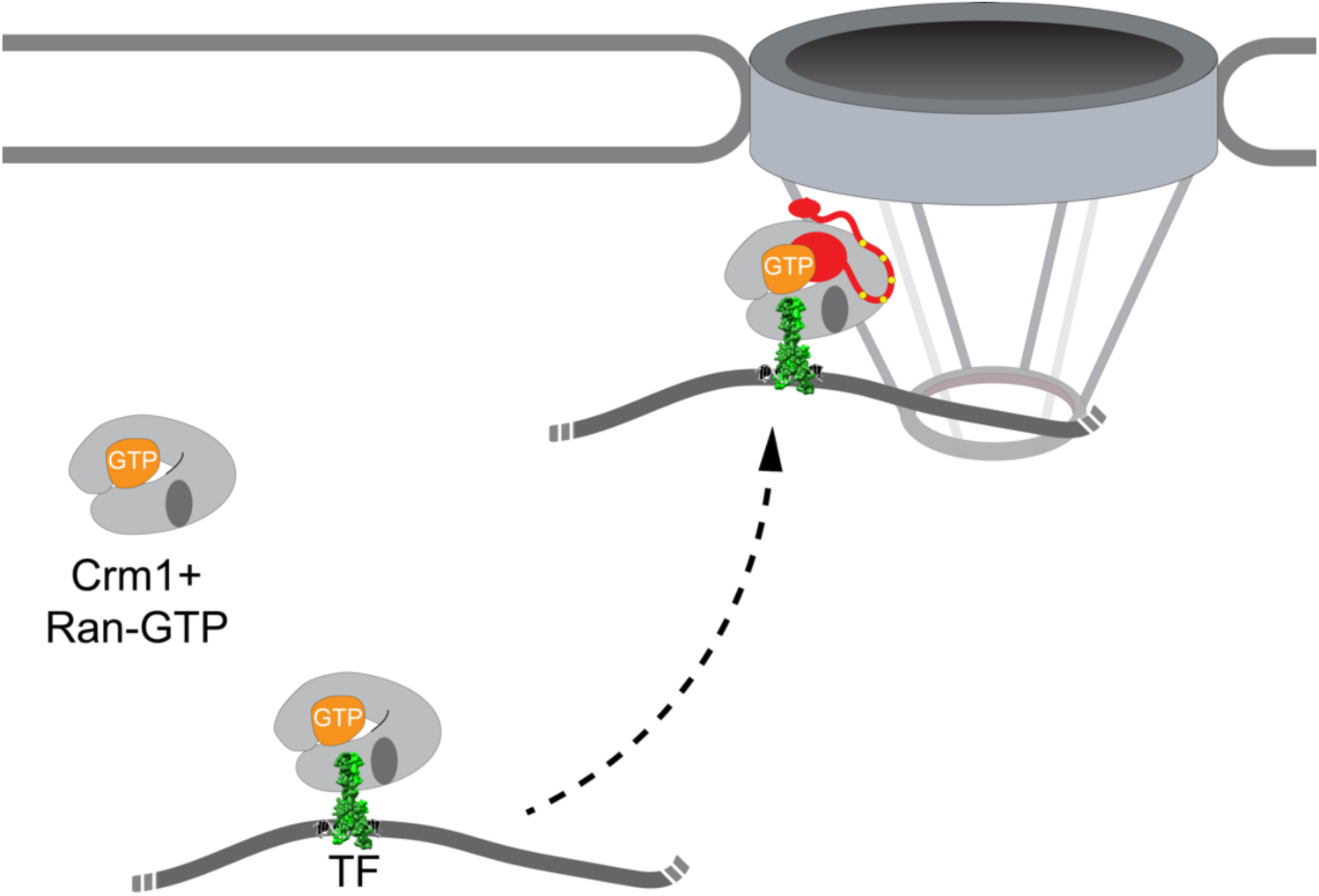
Model for Crm1 function as an adaptor for TF-NPC docking. Crm1 (likely bound to Ran-GTP) associates with TFs bound to DNA in the nucleoplasm. This TF-Crm1 complex undergoes random subdiffusion in the nucleus ^68^. When it encounters the NPC, it interacts with Nup2 (and potentially other Nups) and forms a docking complex.

Inhibiting Crm1 with LMB or depletion by grAID leads to a global loss of Nup2 association and loss of peripheral localization of both Gcn4 targets and non-Gcn4 targets. This suggests that many TFs rely on Crm1 to mediate interaction with the NPC. If so, then other TFs should also have positioning domains that are necessary and sufficient to mediate peripheral localization. While the PD_GCN4_ is highly conserved among Gcn4 homologs from fungi, we have not identified related sequence motifs in other yeast TFs. It is possible that different TFs undergo distinct interactions with Crm1, so they may have functionally equivalent domains with dissimilar sequences. This is akin to the relationship between the amino acid sequence and function of transcriptional activation domains (acidic (Gcn4, VP16), proline-rich (AP-2), glutamine-rich (Sp1), serine/threonine-rich (NFkB)). While such domains lack amino acid sequence identity, they participate in similar interactions with co-activators and Mediator and are sometimes functionally interchangeable ^69^.

Both degradation of Crm1-GFP and LMB treatment of the Crm1-T539C mutant perturb the association of Nup2 with chromatin, suggesting that they both reflect loss of function. LMB inhibits Crm1 by covalently modifying cysteine 539 within the nuclear export binding site ^39^, which specifically blocks binding of nuclear export cargo via its NES. Because the interaction of Crm1 with Gcn4 is not dependent on Ran-GTP, it is surprising that LMB affects targeting to the NPC. There are several possible explanations for this effect. LMB may induce a change in Crm1 localization, depleting if from the nucleus, as is seen in A549 cells ^70^ and *X. laevis* ^71^. Alternatively, LMB may trap Crm1 in a complex that is incompatible with NPC docking because of how it affects interaction with either the TFs or Nups. Finally, it is possible that Crm1 interacts with TFs through the NES binding site in a manner that is not sensitive to Ran-GTP binding. Future work will test these possibilities.

Based on co-occupancy with RNA polymerase II, genetic interactions with Mediator and strong global defects in nascent transcription upon inhibiting Crm1 or Nup2, we conclude that the interaction with these factors promotes transcription of hundreds of yeast genes. However, this defect is largely masked when total mRNA levels were measured, suggesting that it is buffered by reducing mRNA turnover. Such “transcript buffering” has been reported in yeast and mammals in response to global defects in transcription such as loss of histone acetyltransferases, TFIID, or defective alleles of RNAPII ^49,50^. This may explain why the loss of Nup2 has a relatively modest effect on fitness.

Although the effect of Crm1 on transcription was very rapid (i.e., within 30 minutes of LMB treatment), the effect of Nup2 degradation was slow, requiring 4-16h. This suggests that Crm1 has a more direct role than Nup2. Because depletion of Nup2 leads to rapid loss of peripheral localization ^29^, we conclude that Crm1 binding, rather than localization at the nuclear periphery or interaction with the NPC, promotes transcription. Consistent with this notion, Crm1 binds to many sites in the genome that do not interact well with Nups, raising the possibility that these sites may localize in the nucleoplasm but rely on Crm1 for maximal transcription.

How does Crm1 impact transcription? We propose that Crm1 promotes stronger transcription by both strengthening TF binding to DNA to increase TF occupancy, and recruitment of Nups. Nups are mostly unstructured proteins that physically interact with histone modifying enzymes. Thus, Nups may both facilitate histone modifications and enhance the formation of phase-separated condensates such as those frequently associated with transcription. Indeed, tethering Nup153 to chromatin promotes the formation of phase separated foci that include RNAPII and Mediator ^72^. We propose that Nups, together with Mediator, strengthen enhancer function to promote burst frequency. Consistent with this idea, the Gcn4 *pd* mutation leads to a defect in RNAPII recruitment to the promoters of Gcn4 target genes (JV & JHB, manuscript in preparation).

While it is unclear how Nups are recruited to particular sites in metazoans, Crm1 may also serve as an adaptor between TFs and Nups in mammals. In several leukemias, Crm1 co-occupies the *HOX* locus with the leukemogenic NUP98-HOXA9 or SET-NUP214 fusion proteins or the mutant nucleolar protein NPM1c. Both Nup98 and Nup214 are intrinsically disordered Phe-Gly repeat proteins and NPM1c has an unrelated intrinsically disordered domain that promotes phase separation ^73^. Inhibiting Crm1 disrupts binding of the Nups and NPM1c to the *HOX* locus ^74–76^. Likewise, fusion of Crm1 to the histone methyltransferase AFT10/MLLT10 drives leukemia by upregulating the *HOXA/B* gene cluster expression ^77^, presumably because Crm1 is recruited strongly to this locus in stem cells ^75^. Future work will determine if Crm1 serves as a universal, conserved adaptor for TFs to recruit Nups. Also, because Nups are involved in other types of transcriptional regulation (*i.e.*, silencing and epigenetic poising), it will be important to determine Crm1’s role in these phenomena.

## Methods

### Chemicals, media and growth conditions

Unless noted otherwise, all chemicals were from Sigma-Aldrich (St. Louis, MO), restriction and modifying enzymes were from New England Biolabs (Ipswich, MA), DNA oligonucleotides were from Integrated DNA Technologies (Skokie, IL), yeast media components were from Sunrise Science Products (Knoxville, TN). Media were prepared as described ^78^ and yeast cultures were grown at 30°C in synthetic complete glucose (SDC) medium unless indicated otherwise. For peripheral localization experiments, cells were grown at room temperature (RT) in YPD overnight and then shifted to SDC (± histidine or ± histidine) for approximately 1h before imaging. Experiments with Gcn4 target genes were grown in SDC-His, respectively, to induce gene expression. Auxin-Inducible Degron (AID) and <u>gr</u>een targeting AID (grAID) experiments were grown overnight in SDC at RT and then diluted in SDC and grown at 30°C for >4 h before being treated with either 500 µM 3-indole acetic acid (3-IAA; AID) for 1h or 1 µM estradiol + 1 µM 5-phenyl-IAA (5-Ph-IAA; Fisher Scientific) for 2h (grAID) prior to harvesting for ChEC seq. Cultures for SLAM-seq were grown in SDC overnight, diluted in SDC and grown at 30°C for > 4h before switching the cells into SDC-Ura and adding 2 mM 4-thiouracil.

### Yeast strains, plasmids, and molecular biology

All yeast strains were derived from the W303 strains CRY1 (MAT**a** ade2-1 ura3-1 trp1-1 his3-11,15 leu2-3,112 can1-100), CRY2 (*MAT*a *ade2-1 ura3-1 trp1-1 his3-11,15 leu2-3,112 can1-100*; Brickner and Fuller, 1997), or BY4741 (MAT**a** his3Δ1 leu2Δ0 lys2Δ0 ura3Δ0). All strains used in this study are listed in Table S3. Yeast genomic DNA was extracted from strains using the protocol outlined in Looke et al. 2011. PCR purification and gel extraction kits were purchased from Zymo and plasmid purification kits were obtained from Qiagen.

A *pd* mutant Gcn4 strain (JBY536) was generated using CRISPR-Cas9 mediated mutagenesis of the Gcn4-GFP as described ^29^ using the following double stranded repair DNA: 5’-AGTCGTTAAGAAGTCACATCATGTTGGAAAGGATGACGAATCcAGACTaGAcCAcCTAGGTGcTGccGCTgcCAACCGCAAACAGCGTTCGATTCCACTTTCTCCAATTGTG -3’. A wild type strain (JBY537; *GCN4-sm*) with matching silent mutations in the guide sequence was generated using the following double stranded repair DNA: 5’-AGTCGTTAAGAAGTCACATCATGTTGGAAAGGATGACGAATCcAGACTaGAcCAcCTAGGTGTTGTTGCTTACAACCGCAAACAGCGTTCGATTCCACTTTCTCCAATTGTG-3’. After evicting the CRISPR-Cas9 plasmid, mutant strains were confirmed by colony purifying, PCR amplification and DNA sequencing. To generate His-strains, the His5MX marker was replaced in JBY536 and JBY537 by transforming with the KanMX cassette and selecting for G418r, His-, generating JBY550 (*gcn4-pd*) and JBY551 (*GCN4-sm*). *LYS2* was then disrupted in these strains using the *KanMX* to produce JBY557 (*gcn4-pd*) and JBY558 (*GCN4sm*), which were used for the SILAC mass spectrometry experiment.

Leptomycin B-sensitive reporter strain and parental strains with the T539C mutation were a gift from Dr. Eric Weiss (Northwestern University). This strain was transformed with the appropriate LacO, LacI and pER04 membrane marker plasmids to generate strains for chromatin localization experiments ^34^. GFP-tagged proteins in LMB-sensitive strains were made via transformation and homologous recombination of PCR-amplified DNA with 50 bp of homology to the 3’ end of the CDS of the proteins of interest. The GFP was amplified from a gBlock alongside a GSlinker and *Kan^R^* marker. Successful transformations were confirmed by PCR, sequencing around the insert, and imaging. LMB-sensitive were grown overnight in YPD before shifting into SDC ≥1h prior to imaging and treating with 185 nM LMB at 30°C for 30’ prior to imaging.

MNase strains were constructed by amplifying the 3xFLAG-MNase with *Kan^R^* marker from pGZ108 with primers targeting the 3’ end of the CDS of the protein of interest as previously described ^42^. After transforming the PCR construct into cells and selecting with YPD+g418 plates, strains were confirmed by immunoblotting and sequencing. ChEC-seq2 was carried out as described ^45^.

To generate Nup-LexA fusion proteins in strains from the GFP collection, we used a strategy described previously ^29^. The p7-GFPrepl-LexA insert was PCR amplified and transformed into 30 strains having nucleoporin proteins tagged with GFP. Transformants were plated on YPD containing G418, screened for -His as well as imaged by confocal microscopy for the loss of the GFP signal. Proper fusion and expression of selected NPC-LexA DBD proteins was confirmed by PCR and DNA sequencing as well as western blot against LexA DBD. The 27 *MAT***a** Nup-LexA fusion protein strains were crossed against *MAT*α strains expressing LacI-GFP with the nuclear envelope and cortical ER labeled with mCherry (pER04)^34^ and having either p6LacO128-LexABS or p6LacO128 integrated at *URA3*. Diploid strains were selected on SDC -Ade -Ura plates. All diploid strains were visualized on a Leica SP8 (Northwestern Biological Imaging Core Facility). We also generated MNase fusions by replacing the GFP tag from strains in the GFP collection, FLAG-MNase was cloned in place of LexA from p7-GFPrepl-LexA, producing p7-GFPrepl-MNase ^29^. The GFP homology-MNase-*ACT1* 3’UTR – P*_RPL13A_*-*KmR*-*ADH1* 3’UTR - GFP homology was amplified by PCR and transformed into strains having Nup tagged with GFP. Fusions were confirmed by western blot and digestion of genomic DNA upon permeabilization and addition of calcium.

Plasmids pAFS144 ^33^, pFA6a-kanMX6 ^79^, p5LacI-GFP, pER04, pZipKan ^34^, p6LacO128 ^11^, p6LacO128-LexABS ^23,29^, p6LacO128-HIS4 ^23^, pELW749 ^80^, and p7-GFPrepl-LexA ^29^ have been described. The pGEX-WT PD plasmid was generated from pGEX4T-2 by cloning amino acids 205-231 as a BamHI fragment downstream of GST. The PD-DBD was cloned both upstream of GST into pET28a (pET28a-PD-DBD-GST; amino acids 189-281; Figure 6B, C & F) or downstream of GST in pGEX6p-2 (pGEX-TEV-PD-DBD and pGEX-TEV-pdmt-DBD; amino acids 181-281; Figure 6D) because the GST-pdmt-DBD protein was better behaved than the pdmt-DBD-GST protein. The pET28a-GSP1 plasmid was made by cloning a codon-optimized gBLOCK (IDT) of the entire coding sequence of *GSP1* as a *Nco*I + *Hin*dIII into pET28a. The pGEX4T-1-PKI NES plasmid was constructed from oligonucleotides annealed and ligated into pGEX4T-1 digested with *Xho*I + *Bam*HI. The pGEX6p-NUP2C plasmid was made by cloning a codon-optimized gBLOCK (IDT) encoding the final 225 amino acids of *NUP2* as a *Bam*HI + *Xho*I into pGEX6p-2. The pTG020 plasmid was constructed using PCR amplification of ostirF74G from pMK419 ^47^, and of pRSII402 (Addgene #35434). Fragments were purified before mixing with HiFi Assembly Mastermix (NEB) for 30’ at 50°C. Assemblies were transformed into 10-beta competent cells (NEB) and plated onto LB agar plates for overnight incubation at 37°C. The next day, colonies were picked for overnight incubation in liquid LB at 37°C. After ∼18 hours, colonies were purified and sent for whole plasmid sequencing through Primordium Labs. pTG020 digested with StuI was transformed into yeast. pTG023 was constructed using PCR amplification of Plas-gRNA-LEU (Addgene #309094), a gBlock designed to fuse csGBP-mAID (IDT) based on csGBP from Ariotti et al. 2018 and mAID from Yesbolatova et al. 2020. Fragments were assembled and confirmed via sequencing as described above. pTG023 was digested with EcoRV to transform this construct into yeast strains.

### Confocal microscopy

Chromatin localization experiments were carried out as described ^34^ on a Leica SP-8 confocal microscope (Northwestern Biological Imaging Facility). Briefly, *z*-stacks of ≥ 5μm, comprising the whole yeast cell, were collected and gene positioning was scored within the slice with the most focused, most intense LacO/LacI-GFP spot. For experiments in which we scored peripheral localization, ≥30 cells were scored per biological replicate and at least three biological replicates were scored for each strain or condition.

### Stable Isotope Labelling by Amino acids in Cell culture (SILAC) mass spectrometry

WT Gcn4 strain JBY558 was cultured media with 0.1 mg/ml Lysine-^12^C6 ^14^N2 lysine and the *gcn4-pd* mutant strain JBY557 was cultured in media with 0.1 mg/ml ^13^C6 ^15^N2 lysine overnight and then shifted for 1h into 2L SDC-His + heavy or light lysine before harvesting. Cells were harvested by filtration, scraped into a syringe and frozen as noodles. Cells were cryomilled in a Retch MM400 3 x 3min. 5g yeast lysate powder was resuspended on 40ml cold lysis buffer (20mM Hepes-KOH pH 7.4, 50mM KOAc, 20mM NaCl, 0.5% Triton-X100, 0.1% Tween-20, 2 mM MgCl2, 10% glycerol, 1mM DTT + protease inhibitors) by vortexing. Insoluble material was removed by centrifugation at 10,000 x *g*, 10 min and 2mM glutaraldehyde was added to the lysate. After 5 minutes, glutaraldehyde was quenched with 50 mM of lysine. The lysate was added to 50µl sepharose coupled with LaG16 anti-GFP nanobody. Beads were rotated for 1h at 4°C, washed three times in lysis buffer. Affinity purified proteins were eluted with SDS sample buffer and pooled, separated on replicate lanes on a 10% SDS PAGE gels. After staining, each lane was cut into 20 equal size slices, each of which was subjected to quantitative tandem mass spectrometry at the Northwestern University Proteomics Facility.

### Purification of Crm1-x-GFP and Nup2-x-GFP

Anti-GFP pulldown experiments were performed as described, with the following modifications ^55^. After cryomilling, 1g of Crm1-x-GFP or 2g of Nup2-x-GFP powder was resuspended in 8 ml of either Crm1 pulldown buffer (20mM Hepes-KOH pH7.4 250mM Na Citrate 150mM KOAc 0.1% Triton X-100 + protease inhibitors) or Nup2 pulldown buffer (20mM HEPES pH7.4 0.5% Triton X-100 0.1% Tween-20 250mM Na Citrate 150mM NaCl), respectively. The powder was vortexed to resuspend the powder and the lysate was spun 10 min at 11,000rpm in Beckman JLA 16.250 rotor. The supernatant was added to 5mg Dynabeads coupled with LaG94-10 nanobody and incubated 30 minutes at 4°C. The beads were harvested on a magnetic rack, washed three times in the respective pulldown buffer and resuspended in 20µl 5% SDS 500mM NH_4_OH and heated at 70°C, 5 minutes. The beads were removed on a magnetic rack and the supernatant was collected. 5µl of the eluate was run on a 4-12% NuPAGE gel and the remainder was dried in a Speedvac for 2h, with heating.

Samples were processed for mass spectrometry as described ^81,82^ with the following modifications: peptides were generated from proteins using S-Traps (ProtiFi, Fairport NY) according to the manufacturer. Peptides were then purified over a C18 StageTips (Pierce) and analyzed by LC-MS using an Orbitrap Exploris mass spectrometer coupled with an Easy-nLC system (Thermo Fisher Scientific). SpectroMine software (Biognosys AG) was used for label-free quantitation (LFQ). To compare LFQs across biological replicates, proteins were normalized to the antigen.

### ChEC seq2 and SLAM-seq and data analysis

ChEC-seq2 and data analysis in R were performed as described ^45^. For the grAID experiment (Figure 3), cleavage was carried out in SDC + digitonin + 2mM calcium. SLAM-seq was performed as described ^48,83^. Yeast cells were grown in synthetic complete medium containing glucose (SDC) before Crm1 was inhibited by treating the cultures with 185 nM leptomycin B (LMB) in ethanol for 30 min at 30°C. Nup2 was degraded by treating cultures with 500 uM indole-3-acetic acid (3-IAA) in ethanol for 16 hours at 30°C. Following treatment, cultures were treated with 0.2 mM 4sU, for 6 min at 30°C. Cells were spun down and snap frozen. Total RNA was extracted using Phenol:Chloroform:IAA (25:24:1) extraction. 5 ug total RNA was treated with 10 mM iodoacetamide for 15 min at 50C. 3’ mRNA-seq libraries were made using the QuantSeq 3’mRNA-seq Library Prep Kit for Illumina (FWD; Lexogen) following manufacturer’s recommendations and previously published protocol. All experiments had at least three biological replicates. SLAM-seq data was analyzed by SLAMDunk ^48^ and DESeq2 ^84^ was used to identify mRNAs and nascent transcripts that changed significantly.

### grAID (<u>gr</u>een fluorescent protein-targeted AID)

grAID strains were grown in SDC overnight at RT or 30°C before diluting in SDC the next day and growing at 30 ° C for > 4h. Cells were then treated with 1 µM estradiol (E2, Sigma Aldrich) for 30 minutes before treating with 1 µM of 5-Ph-IAA (Fisher Scientific) for 2h and harvesting for immunoblotting as well as ChEC-seq2.

### Protein purification

H_6_-Crm1 was expressed in BL21(DE3) and purified as described ^80^. H_6_-GSP1, GST-NES, GST-NUP2C, GST, GST-DBD, GST-PD, PD-DBD-GST and GST-PD-DBD were expressed in BL21(DE3) in LB + appropriate (either ampicillin or kanamycin) antibiotic (2L) at 37°C for 3h, harvested, washed in water and then resuspended in 50ml lysis buffer (20mM Tris pH 7.5 400mM NaCl 0.5mM EDTA 2mM DTT and protease inhibitors) and frozen in 2 x 25ml tubes in liquid N_2_. Cells were thawed and lysed in an Avestin C3 high pressure homogenizer (4 x 15,000 psi). Nucleic acids were precipitated by addition of 1ml 5% polyethylene amine while mixing. Insoluble material was removed by centrifugation 30-60 minutes at 40,000 rpm in a Beckman TLS55. To the supernatant, NaCl was added to 1M and passed over a 5ml GSTrap column using a peristaltic pump. The column was washed 1 x 10ml lysis buffer + 1M NaCl and then transferred to an AKTA Purifier FPLC, where it was washed with 10ml 20mM Tris pH7.5, 100mM NaCl, 0.5mM EDTA, 2mM DTT and then eluted over a 0-100% gradient of 20mM Tris pH7.5, 40mM glutathione, 100mM NaCl, 0.5mM EDTA, 2mM DTT. Fractions were analyzed by SDS PAGE, and peak fractions were pooled and dialyzed overnight into storage buffer (20mM Tris pH 7.5, 300mM NaCl, 0.5mM EDTA, 2mM DTT). For experiments in which tags were removed from Gcn4 or Nup2C, the peak fractions were incubated with Prescission protease overnight during dialysis into 20mM Tris pH7.5, 150mM NaCl, 0.5mM EDTA, 2mM DTT. The Gcn4 PD-DBD (aa 181-281) fragment was bound to a 5 ml HiTrap S column, washed with 10ml 20mM Tris pH7.5, 100mM NaCl, 0.5mM EDTA, 2mM DTT, and eluted with a gradient of 0-100% 10ml 20mM Tris pH7.5, 2.5M NaCl, 0.5mM EDTA, 2mM DTT, pooled peak fractions and dialyzed into 20mM Tris pH7.5, 100mM NaCl, 0.5mM EDTA, 2mM DTT overnight at 4°C. Glycerol was added to 5% and aliquots were frozen in liquid N_2_.

Ran was loaded with nucleotide by combining 100µM H_6_-Gsp1 with 20mM GDP or GTP and incubating at 30°C for 20 minutes. For reactions involving Ran, it was diluted to 10µM, with 2mM nucleotide.

### Fluorescence Polarization

For Gcn4 binding experiment, 110µl of 40µM Gcn4 PD-DBD was labeled using the AlexaFluor 488 labeling kit (ThermoFisher) according to the manufacturer’s instructions. The reaction was quenched by addition of 900µl 100mM Tris pH7.5 and purified using a 3000 MWCO concentrator spin filter. For the Nup2C experiments, GST-NUP2C was purified and dialyzed into 0.3M sodium bicarbonate pH 9.5 overnight at 4°C, aliquoted and frozed in liquid nitrogen.10mg of GST-Nup2C was combined with Atto488 labeling mix and incubated 2h at 4°C. Labeled protein was separated from unlabeled protein using a PD-10 gel filtration column and stored at 4°C.

### Electrophoretic Mobility Shift Assay

In 20µl reaction containing 20% glycerol, 100mM KCl, 20mM HEPES pH 6.8, 0.2mM EDTA, and 0.042% bromophenol blue, 200nM GST-Gcn4 PD-DBD (amino acids 181-281) was combined with 5pmoles AlexaFluor488-labeled Gcn4 binding site + 50µg poly dIdC ± 1µM Crm1 ± 2.5µM Ran-GTP. The reaction was incubated for 15 minutes at room temperature and then separated on a 6% DNA retardation gel in 0.5X TBE running buffer (ThermoFisher). The gel was imaged with a Sapphire^TM^ multimode imager in the Keck Biophysics facility at Northwestern University.

## Acknowledgements

The authors thank Professor Eric Weiss (Northwestern University) for sharing the leptomycin B-sensitive yeast strain, Professor Reza Vafabakhsh (Northwestern University) for sharing nanobody-coupled beads, Alexis Jacob for help with microscopy, Dr. Trevor Van Eeuwen (Rockefeller University) for help with figures and members of the Brickner laboratory for helpful comments on the manuscript. This work was supported by NIH grants R35GM136419 (JHB), P41 GM109824 (MPR & BC), R01 GM112108 (MPR), T32 NIGMS GM008061 (TG & DJV), F32 GM153164 (CC) and a predoctoral fellowship from the National Science Foundation (DJV).

## Primary data

Sequencing data (fastq files for each replicate) associated with this project were submitted to the National Library of Medicine under the following accession numbers: PRJNA1109451, PRJNA1109452, PRJNA1109460.

---

**Figure S1, related to Figure 1.**
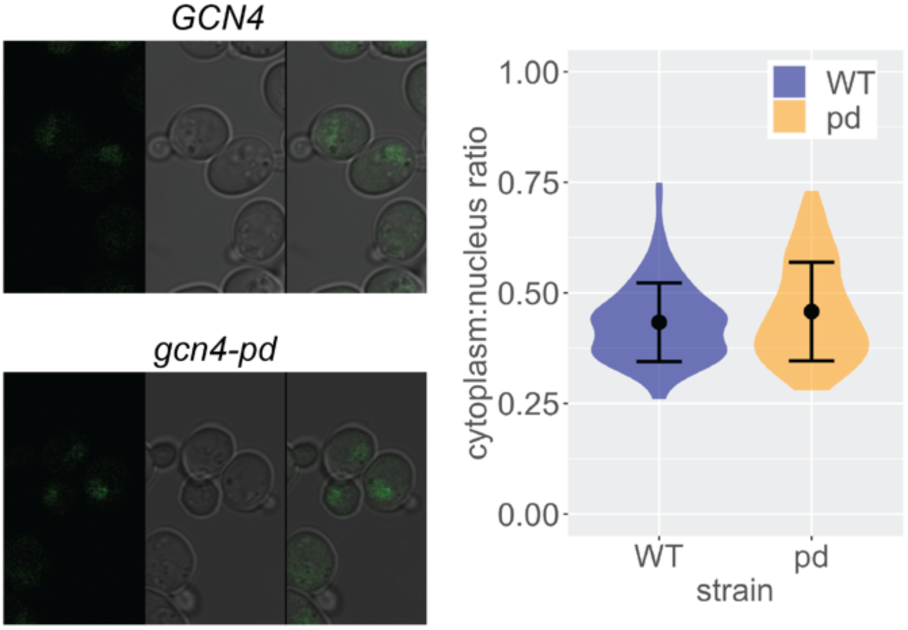
The pd mutation does not alter Gcn4-GFP localization. Left: confocal micrographs of *GCN4-GFP* and *gcn4-pd-GFP* strains starved for 1h for histidine. Right: the fluorescence intensity for comparable areas of the nucleus and cytoplasm was measured in 30 cells. The mean ratio of fluorescence of cytoplasm to nucleus is plotted ± standard error of the mean.

**Figure S2, related to Figure 2.**
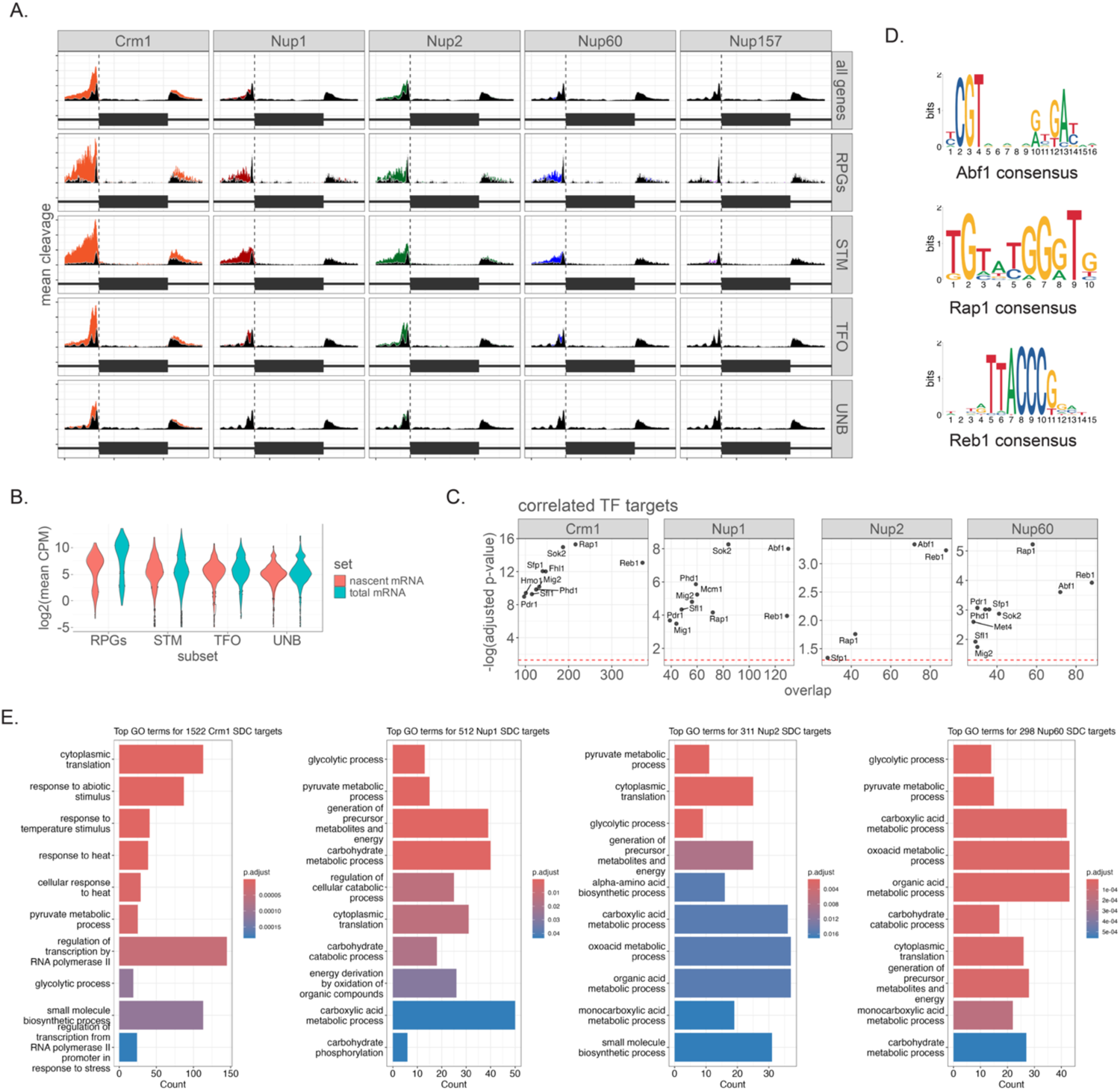
Crm1 and Nups associate with yeast enhancers of strongly expressed genes. **A.** Mean cleavage by Crm1, Nup1, Nup2, Nup60 and Nup157 over normalized length metagene plots flanked by 700bp upstream and downstream of different categories of genes ^44^. **B**. Total and nascent RNA counts per million reads from CRY1 grown in SDC for the subsets used in Figure 2 ^44^. **B.** Top enriched TF targets for each protein. The number of genes that overlap between those identified by peaks of Crm1 and Nups with the indicated TFs was plotted against the Bonferroni-adjusted p-value (Fisher’s Exact test). **C**. Consensus motifs for Abf1, Rap1 and Reb1 from https://jaspar.elixir.no/. **D**. Top 10 Gene Ontology terms enriched for genes adjacent to high-confidence sites for each protein, with the number of genes and the adjusted p-value for each term.

**Figure S3, related to Figure 3.**
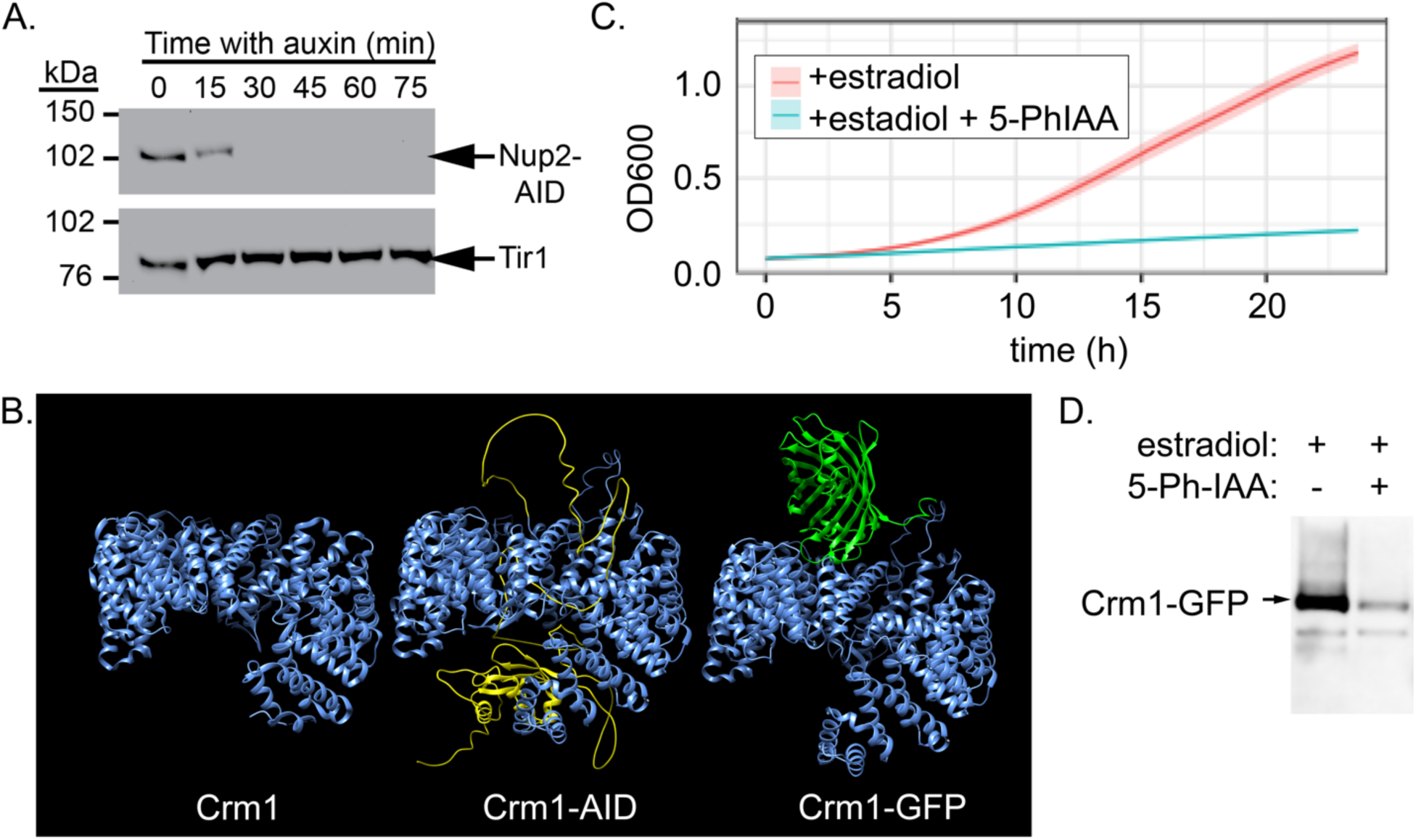
**A.** Nup2 auxin-induced degradation. Protein was extracted from cells expressing Nup2-myc-IAA7 and Tir1-myc at the indicated times after addition of 0.5mM IAA, separated by SDS PAGE, transferred to nitrocellulose and immunoblotted for the myc tag. **B**. Alphafold2 predictions for Crm1, Crm1-AID and Crm1-GFP. **C**. growth of Crm1-grAID strains in the presence of estradiol ± 5 Ph-IAA. **D**. Protein levels of Crm1-GFP 2h after addition of estradiol ± 5-Ph-IAA.

**Figure S4.**
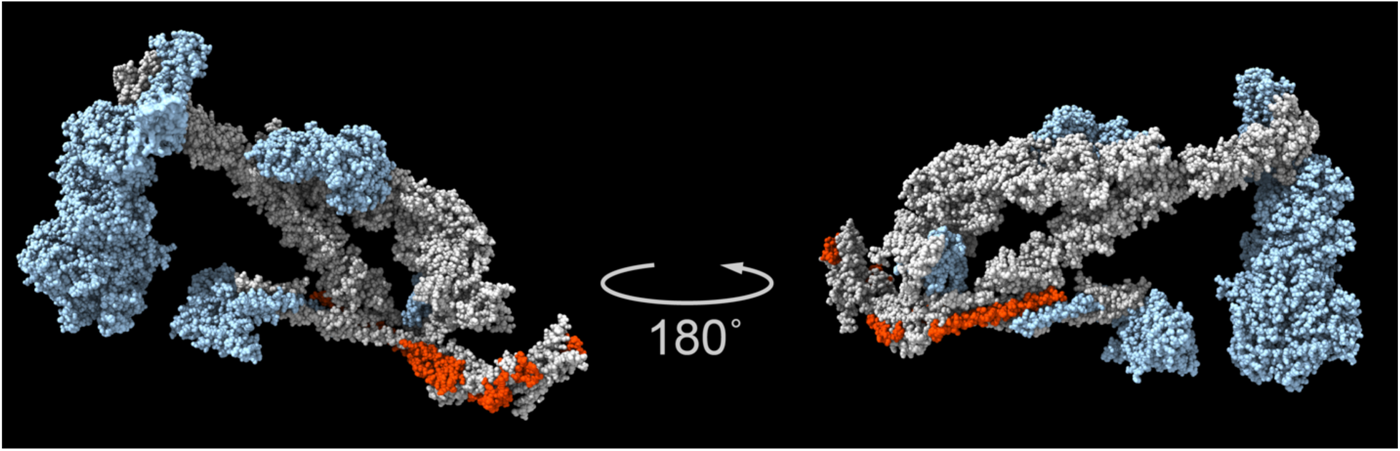

**Figure S5, related to Figure 5.**
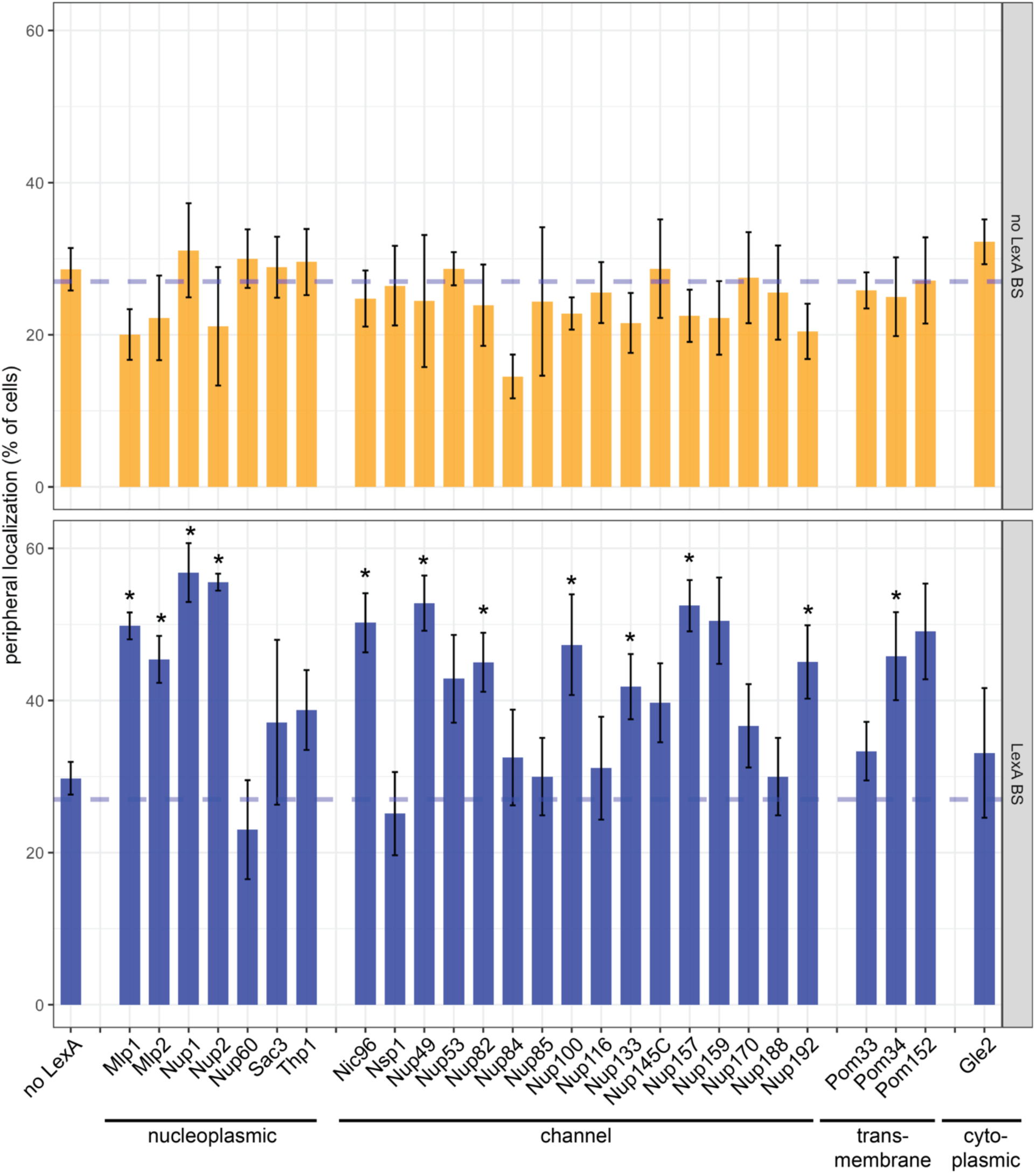
Peripheral localization mediated by LexA tethering of chromatin to the NPC. The LexA DNA binding domain was fused to the carboxyl terminus of each of the indicated nuclear pore proteins. These strains were crossed against a strain expressing GFP-LacI and PHO88-mCherry and having a LacO array integrated at *URA3*. One version of this strain possessed the LexA binding site at *URA3* (bottom panel) and the other did not (top panel). Peripheral localization of *URA3* was measured in three biological replicates of ≥ 30 cells each. The mean peripheral localization (% of cells) ± the standard error of the mean is plotted. The dashed blue line represents the expected colocalization with the nuclear periphery for a randomly positioned locus ^11^.

**Figure S6, related to Figure 6.**
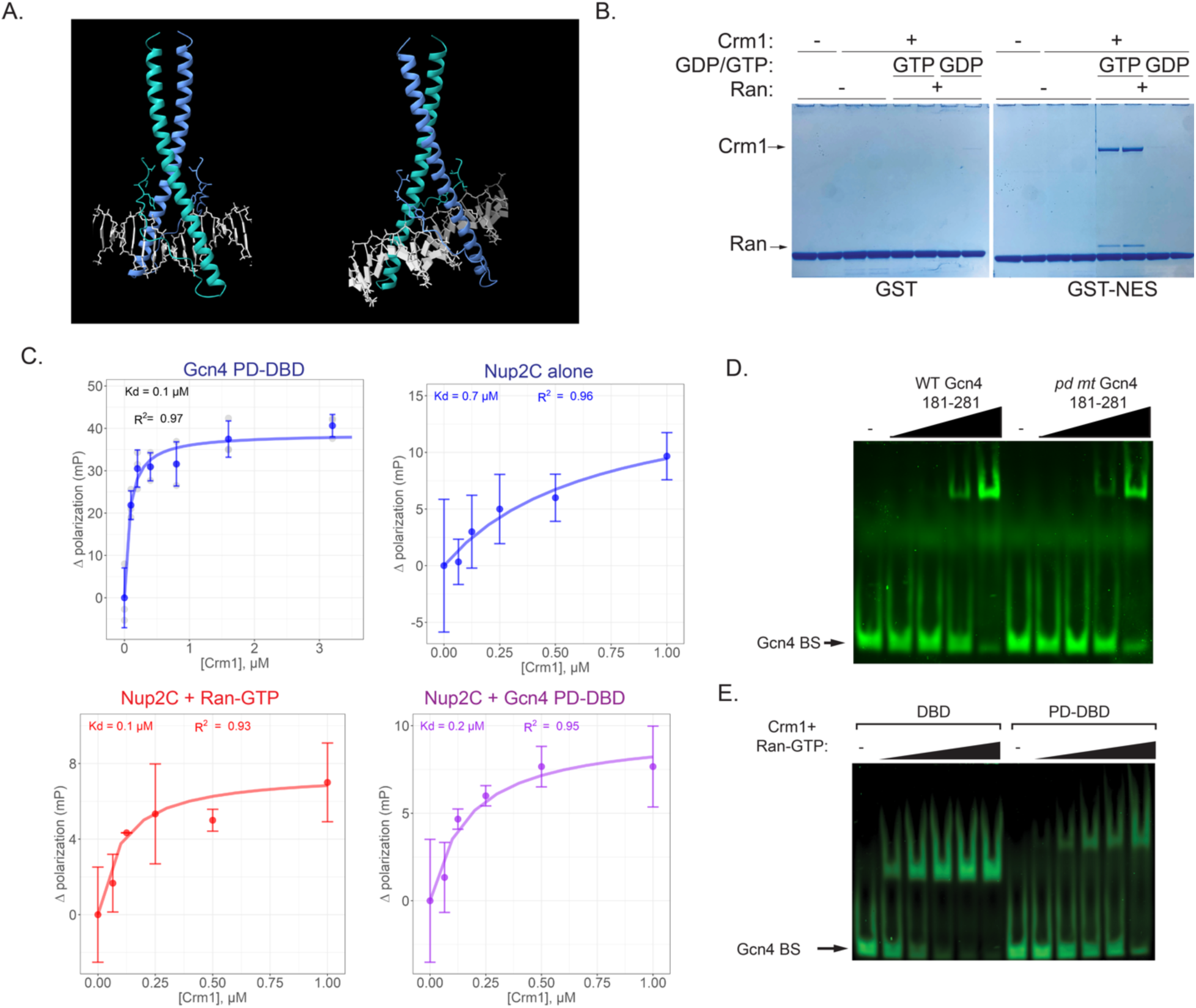
Biochemical characterization of Crm1 interactions. **A**. Alphafold3 prediction of the structure of Gcn4 PD-DBD (amino acids 181-281), bound to DNA from two different perspectives. The side chains of the PD amino acids (VVAY) that are sensitive to mutations are shown. **B**. Biochemical reconstitution of the nuclear export complex. GST-NES (from PKI) was bound to magnetic glutathione beads, washed and the beads were incubated with Crm1, Ran-GDP and/or Ran-GTP as indicated. Crm1 and Ran binding was only observed in the presence of Ran-GTP. **C & D**. Fluorescence polarization with 40pM AlexaFluor488-labeled Gcn4 PD-DBD (untagged; panel **C**) or 25pM Atto488-labeled Nup2C (untagged; panel **D**) was measured in the presence of the indicated concentrations of Crm1. In panel **D**, 2µM Ran-GTP or Gcn4 PD-DBD was added (middle and right panels). Curves were fit to generate the indicated Kd values. **E**. EMSA of the Gcn4 binding site incubated with either buffer (-lanes) or the following concentrations of untagged wild type or *pd* mutant Gcn4 PD-DBD (amino acids 181-281): 40nM, 200nM, 800nM and 4µM.

